# Dyr726, a brain-penetrant inhibitor of PI3Kα, Type III receptor tyrosine kinases, and WNT signaling

**DOI:** 10.1101/2025.03.26.645490

**Authors:** Vasudha Tandon, Alessandra Fistrovich, Joaquina Nogales, Febe Ferro, Samantha N Rokey, Carly Cabel, Amy Dunne Miller, Mary Yagel, Christina Duncan, Aditi Atmasidha, Ira Sharma, Gerrit Wilms, Karin Williams, Richard Elliott, Timothy Chavez, Yeng Shaw, Aidan McMahon, Sean Ginn, L. Emilia Basantes, Nathan Bedard, Vijay Gokhale, Nathan Ellis, Alan R Prescott, Stuart J Smith, Ruman Rahman, Walter Becker, Kevin D Read, Anthony J Chalmers, Neil Carragher, Glenn R Masson, William Montfort, Curtis Thorne, Christopher Hulme, Sourav Banerjee

## Abstract

The vast majority of clinical small molecule multi-kinase inhibitors (mKI) report abject failures in targeting cancers with high stem cell contents like high-grade glioma and colorectal cancers. The FDA-approved mKIs to date ablate receptor tyrosine kinase signaling but do not target the paradoxical WNT signaling which is a key survival driver for the self-renewing cancer stem cells. The WNT pathway enhances cancer plasticity and triggers relapse of highly heterogenous tumours. Using *de novo* synthesis and structure-activity-relationship (SAR) studies with blood-brain-barrier (BBB) penetrant mKI scaffolds, we designed a highly potent and selective small molecule inhibitor of PI3Kα, PDGFR/KIT, and the WNT pathway denoted Dyr726. Dyr726 is superior to clinical mKIs and inhibits PI3K-AKT-mTOR and WNT-pathway signaling at multiple nodes thereby impeding proliferation, invasion, and tumour growth. Phospho-proteomic, structural, and target engagement analyses, combined with *in vitro*, *in vivo* efficacy, and pharmacokinetic studies reveal that Dyr726 is a brain-penetrant small molecule which effectively reduces tumour volume and extends survival of murine orthotopic models. Our current work establishes a first-in-class brain penetrant small molecule mKI which simultaneously antagonize the PI3K-AKT-mTOR and WNT pathways in preclinical cancer stem cell cultures, adult and pediatric primary organoids, and orthotopic murine models with positive efficacy in combination with clinical standard of care.

## Introduction

Kinase inhibitors remain one of the most successful classes of small molecules that clinically alleviate cancer^1^. Although only a small fraction of the 540 kinases have been targeted clinically to date, the repertoire of kinase inhibitors expands every year with over 80 molecules now approved by the FDA^2^. However, most kinase inhibitors that are repurposed to target cancers with highly heterogenous microenvironments fail clinical trials. This is partly due to their inability to target the large population of cancer stem cells (CSC)^3^. CSCs’ ability to self-renew and differentiate allows them to reinitiate tumours and maintain tumour heterogeneity leading to therapeutic resistance and relapse^4^; which consequently causes the abject failure of kinase inhibitors in targeting adult and pediatric high-grade gliomas and β-catenin/APC-mutated solid tumours^5, 6^.

Aberrations in RTK signaling pathways have been observed across most cancers^1, 6, 7^. Specifically, in high-grade gliomas, EGFR and PDGFRA/KIT mutations and amplifications are commonplace often compounded with gain-of-function mutations in oncogenic/tumour promoting lipid kinase PI3Kα and tumour suppressor PTEN loss^8–10^. Unfortunately, targeting either EGFR or PI3K isoforms has not been successful with gefitinib, erlotinib, and pan-PI3K buparlisib exhibiting modest to no efficacy along with significant toxicities in brain tumour patients^6^. Targeting type III RTKs PDGFR/KIT with avapritinib on the other hand has shown some exceptional responders in pediatric diffused intrinsic pontine glioma patients^11^. This could be due to the redundancy of the PDGFRA/KIT in regulating the WNT signaling and glioma stemness^12, 13^. The WNT signalling pathway drives cancer stemness, yet there are no approved drugs targeting the WNT pathway currently in the clinic. Previous attempts to directly target WNT drivers proved too toxic^13^. Multiple attempts are underway pre-clinically to generate WNT pathway inhibitors by targeting specific kinases that regulate the β-catenin destruction complex or the splicing of the WNT target genes^14, 15^. PDGFR/KIT have been reported to be the key targets of a novel multi-kinase inhibitor WNTinib which ablates WNT signaling through KIT inhibition in β-catenin-mutated hepatocellular carcinoma^12^. Another small molecule cirtuvivint is currently in clinical trials for advanced solid tumours which causes WNT pathway ablation through inhibition of a class of CMGC kinases called DYRK and CLK^14^. DYRK1A-4 and CLK1-4 have been reported to modulate alternative mRNA splicing for WNT target genes and hence inhibition of the group leads to ablation of the WNT signaling pathway^14, 15^. In fact, our previous works have shown that small molecules targeting the DYRKs have potent *in vivo* efficacy against triple-negative breast cancer and multiple myeloma^16, 17^. Currently, the CDK4/6 inhibitor abemaciclib has been shown to possess potent off-target inhibitory effects on pan-DYRK and pan-CLK, although, its effect on the WNT pathway has not been rigorously evaluated^18^. Abemaciclib is a blood-brain-barrier penetrant inhibitor that is FDA approved for HR+/HER2-breast cancer and is currently being evaluated in glioma clinical trials in combination with standard of care (NCT02977780 and NCT02981940)^19^. Hence, in conjunction with RTK inhibition, WNT pathway targeting could indeed be a key cog in the therapeutic targeting of cancers with high stemness like gliomas and those with β-catenin/APC-mutated solid tumours.

To target multiple oncogenic drivers simultaneously, a recent paradigm shift has occurred in developing kinase inhibitors with pleiotropic, multi-targeted profiles^20^. We have thus developed a single brain-penetrant molecule which can simultaneously ablate PI3Kα signaling, PDGFR/KIT signaling, and the WNT pathway. Using *de novo* synthetic strategies, we generated Dyr726, a potent PI3Kα, type III RTK, pan-DYRK1A-3, pan-CLK1-3 inhibitor. Dyr726 is a first-in-class, brain penetrant, small molecule which can potently target proliferation of primary CSC and organoid cultures, ablate WNT pathway, impede tumour progression, and extend survival of orthotopic *in vivo* model systems.

## Results

### Dyr726 is an efficacious multi-kinase inhibitor

Dyr726 is a potent multi-targeted ATP-competitive Type 1 kinase inhibitor designed to target PI3Kα, type III receptor tyrosine kinases PDGFR/KIT/FLT3, and the DYRK/CLK class of CMGC kinases. Multiple kinase specificity screens clearly confirm the kinome engaged by Dyr726 (Fig 1A; Supplementary Table S1). The molecule did exhibit off-target engagement with Haspin and BRAF (Supplementary Table S1), however, no clear evidence of BRAF cellular target engagement was observed (Supplementary Fig S1). To benchmark the anti-cancer potency, we studied the ability of Dyr726 to impair cellular viability in a panel of genetically diverse adult and pediatric patient-derived primary glioma cells (Fig 1B). Glioma was the ideal cancer-type to assess efficacy of Dyr726 since glioma exhibits heterogenous genetic aberrations in RTKs with very high incidence of dysregulated PI3Kα-mTOR signaling due to frequent PTEN-loss. Furthermore, glioma exhibits stemness with highly plastic and self-renewing subtypes of pro-neural, mesenchymal, and classical glioma stem cells (GSCs) which contribute to the heterogenous, immune-suppressive, and drug-resistant tumour microenvironment. Dyr726 has potent cytotoxic effects on all primary glioma lines tested regardless of genetic drivers and exhibited an EC_50_ of 0.3-3 μM across the cultures (Fig 1B). Next, we assayed the ability of Dyr726 to reduce 3D invasion of glioma cells. Pseudo-glioma U87-MG and primary patient derived glioma GBM102 cells were treated with 500nM Dyr726 which over 4 days significantly reduced invasion of the spheroids into the surrounding matrix (Fig 1C). To benchmark Dyr726 against FDA approved kinase inhibitors, we treated two adult primary glioma neurosphere cultures GBM12 and GBM76 with increasing concentrations of Dyr726 or crenolanib (Type III RTK inhibitor) or avapritinib (Type III RTK inhibitor) or alpelisib (PI3Kα inhibitor) or capivasertib (AKT inhibitor) or abemaciclib (CMGC kinase inhibitor). Compared to the FDA approved kinase inhibitors, Dyr726 was found to be the most potent at reducing the glioma neurospheres in both cultures (Fig 1D). Interestingly, Dyr726 exhibited additive cytotoxicity in combination with 50 μM temozolomide (Fig 1E) and radiation (Fig 1F) which are standard-of-care for GBM. GBM cells treated with increasing dose of radiation in presence of increasing concentrations of Dyr726 exhibited fewer surviving colonies (Fig 1F).

**Figure 1:**
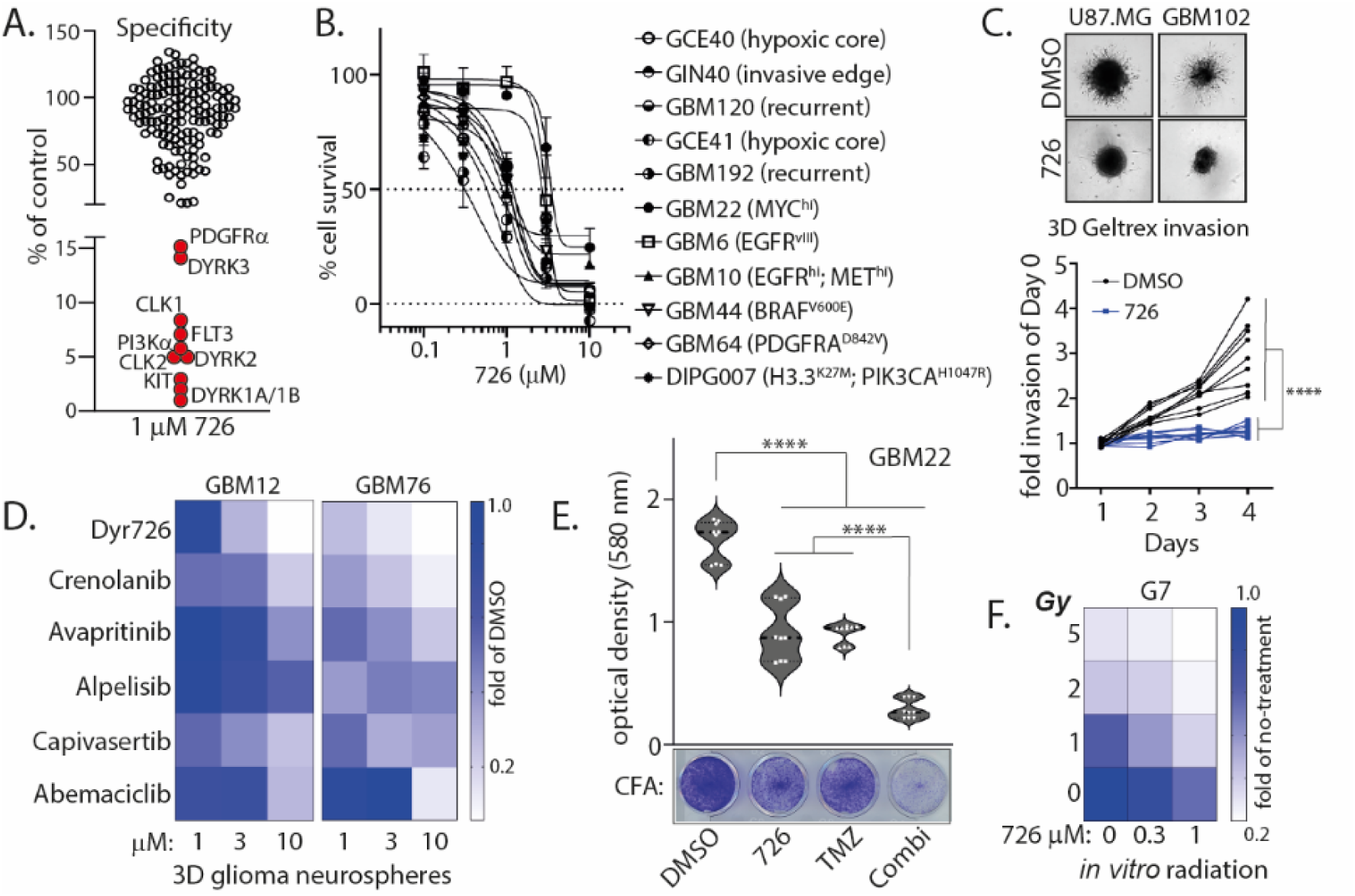
Dyr726 is an efficacious multi-kinase inhibitor (A) Kinase profiling of at 1 μM was carried out at the International Centre for Protein Kinase Profiling (http://www.kinase-screen.mrc.ac.uk/). Please also refer to Supplementary Table T1 and Fig S1 for extended data on kinase selectivity using DiscoverX platform. (B) Dyr726 potently kills a panel of genetically diverse primary patient-derived adult and pediatric glioma cells between 0.3-3 µM EC_50_. (C) 2,000 U87-MG or GBM102 cells were seeded per well with their respective cell culture media in a U-Shaped-Bottom Microplate for 2-3 days to allow spheroids to form. Once spheroids had formed, they were isolated and embedded into Geltrex™. Complete medium was placed on top with either DMSO or 500nM Dyr726. Images were taken after 3 days on the EVOS XL core microscope. The areas of invasion were quantified using FIJI. **** P < 0.0001 2-way ANOVA, mean +/-SD from n = biological replicates. (D) GBM12 and GBM76 neurospheres were allowed to form for 14d and 7d respectively. Once formed neurospheres were treated with Dyr726, crenolanib, avapritinib, alpelisib, capivasertib, abemaciclib, or DMSO control for 14d with the indicated concentrations. After further 14d, neurosphere diameters were measured using FIJI. (E) Colony formation assay was carried out with MGMT methylated GBM22 cells with or without the treatment with 1 μM Dyr726, 50 μM temozolomide (TMZ), and combination (Combi) of both over 7 d. Cells were fixed with 100% EtOH and stained using crystal violet solution. 24h later, the crystal violet solution was reconstituted using 10% acetic acid. Absorbance was measured at 590nm using the TECAN plate reader. (F) G7 cells were pretreated with Dyr726 or DMSO control for 1hr prior to irradiation at the indicated doses. 14d post irradiation at the indicated Gy doses, colonies were fixed and stained and manually counted using FIJI.

### Dyr726 is an isoform specific PI3K**α** inhibitor

Our structure-activity-relationship studies of over 500 molecules produced Dyr726, which is an isoform selective inhibitor for PI3Kα. Specifically, biochemical IC_50_ analysis reveals that Dyr726 is a PI3Kα inhibitor (IC_50_ 3.6nM). It exhibits over a 1,000-fold higher IC_50_ against the β and γ-PI3K isoforms, with no observed inhibition of AKT1 (Fig 2A). Dyr726 also inhibits common PI3Kα activating mutants, E542K, E545K, and H1047R with IC_50_s of 5nM, 3.8nM, and 15.7nM respectively (Fig 2A). In primary patient derived GBM6 cells, Dyr726 inhibits the oncogenic PI3K-AKT-mTOR signaling pathway with complete ablation of phospho-AKT, phospho-PRAS40, and phospho-S6K within 1 h upon titration with Dyr726 (Fig 2B). Indeed, increasing doses of Dyr726 led to marked death in both adult and pediatric high-grade glioma neurospheres. Adult primary GBM215 neurosphere cultures are EGFR amplified and allowed a 45% reduction in neurosphere size upon treatment of Dyr726 (Fig 2C). Similarly, HSJD-DIPG-007 is a H3.3 K27M mutated, PDGFRA amplified, and PI3Kα H1047R mutated pediatric diffused midline glioma neurosphere culture which also exhibited very potent death upon titration with Dyr726 (Fig 2D). To look at global phosphorylation changes upon Dyr726 treatment, three different primary GBM cells GBM6, GBM22, and GIN28 were treated with and without 10 µM Dyr726 for 2 h and quantitative tandem-mass-tag labelled 6-plex phospho-proteomics analysis was carried out. Indeed, pT308 AKT1 was downregulated significantly (Fig 2E). Gene ontology analysis shows that overall, the mTOR signaling pathway is reduced with Dyr726 treatment. However, no changes in MAPK-signaling or phospho-ERK levels were observed. Indeed, phospho-ERK levels are not affected by Dyr726 in EGFR-amplified GBM143 cells treated with human epidermal growth factor (Fig 2F). The MAPK-pathway in GBM143 cells is likely driven by upstream EGFR. Thus, ablation of phospho-AKT with no effect on phospho-ERK activity confirms that EGFR is not inhibited by Dyr726 and the phospho-AKT ablation is due to PI3Kα inhibition alone. Thus, Dyr726 is a highly potent isoform specific inhibitor of PI3Kα.

**Figure 2:**
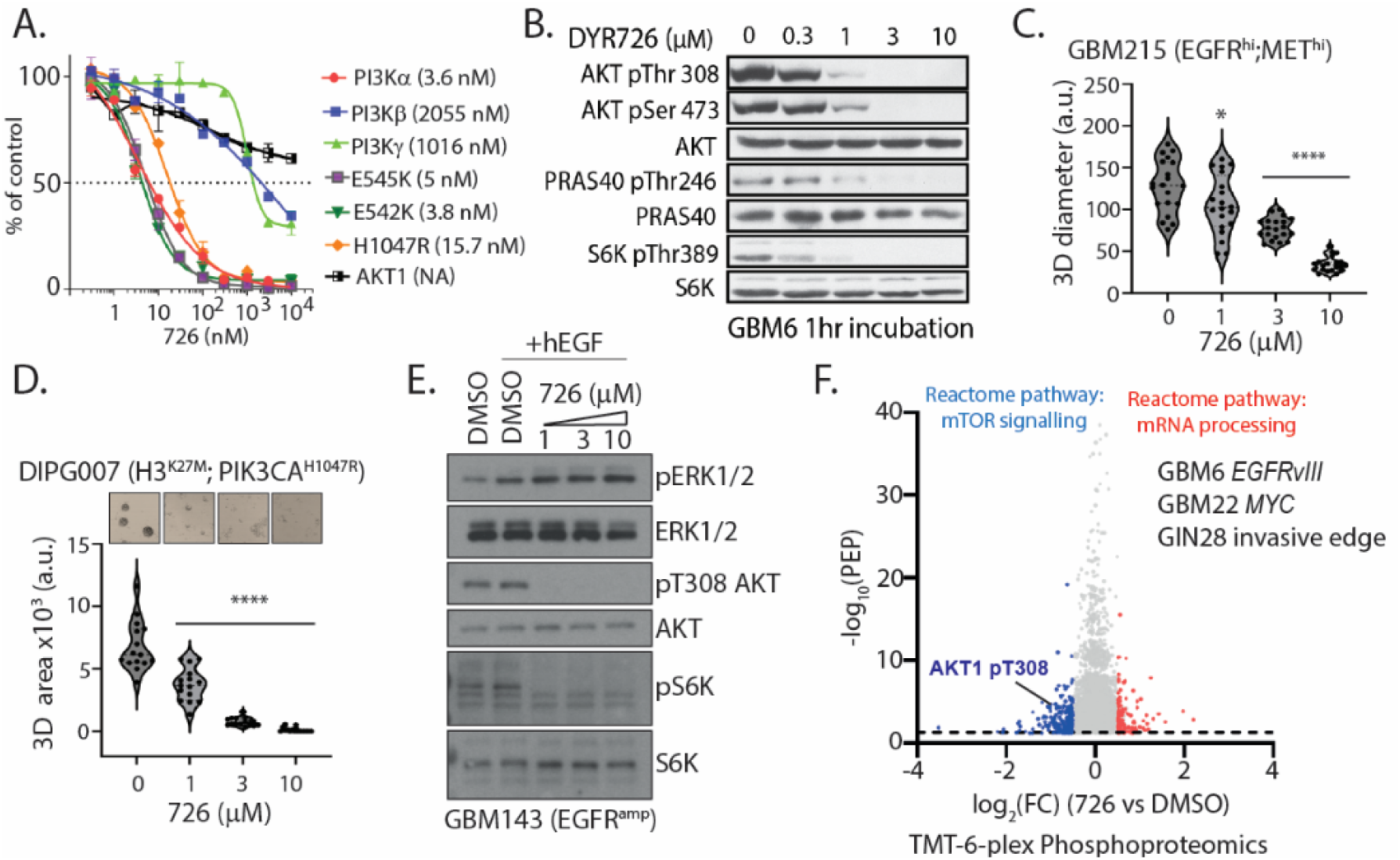
Dyr726 specifically inhibits PI3Kα and gain-of-function mutants. (A) PI3Kα, PI3Kβ, PI3Kγ, PI3Kα E545K, PI3Kα E542K, PI3Kα H1047R, and AKT1 were assayed using the indicated concentrations of Dyr726. The IC_50_ graph was plotted using GraphPad Prism software. (B) GBM6 cells were treated with increasing concentrations of Dyr726 for 1hr. Cells were lysed, and immunoblotting was performed using the indicated antibodies. (C) GBM215 cells were plated at 10,000 cells per well and were incubated for 7d to form neurospheres. Once neurospheres were formed, they were treated with indicated concentrations of Dyr726 for 14d. FIJI was used to measure the diameter of the neurospheres. * P < 0.05; **** P < 0.0001 one-way ANOVA, mean +/- SD from n = biological replicates. (D) HSJD-DIPG-007 cells were plated at 10,000 cells per well and were incubated for 1wk to form neurospheres. Once neurospheres were formed, they were treated with indicated concentrations of Dyr726 for 2wks. FIJI was used to measure the area of the neurospheres. **** P < 0.0001 one-way ANOVA, mean +/- SD from n = biological replicates. (E) EGFR amplified GBM143 cells were treated with hEGF and the indicated concentration of Dyr726 or DMSO control for 2hr. Cells were then lysed and immunoblotting was performed using the indicated antibodies. (F) Volcano plot depicting the log2 of fold phosphorylation change versus -log10(p-value) for phosphosites on secretory pathway proteins. The phospho-peptides marked in blue displayed a statistically significant increase while those in red displayed decrease in GBM6, GBM22, and GIN28 cells treated with 10 µM Dyr726 for 0.5hr. TMT-6-plex phospho-proteomics was carried out using three biologically distinct primary glioblastoma cultures.

### Dyr726 is a type III RTK signaling inhibitor

Although Dyr726 does not inhibit EGFR in cell or biochemically, Dyr726 is a very potent inhibitor of the type III RTKs PDGFRA/B, KIT and FLT3. Biochemical IC_50_ analyses show that Dyr726 inhibits PDGFRA at 62nM, PDGFRB at 76nM, KIT at 13nM, and FLT3 at 55nM (Fig 3A). To benchmark Dyr726 against FDA approved type III RTK inhibitors, we stably overexpressed c-KIT in GBM6 (Fig 3B) and PDGFRA in GBM22 (Fig 3C) cells and activated the KIT and PDGFR-mediated signaling using SCF and PDGF-BB respectively. Both Dyr726 and avapritinib ablated phospho-AKT, phospho-S6K, and phospho-ERK signals although crenolanib exhibited a relapse of the signaling at higher concentrations (Fig 3C). All 3 inhibitors ablated the tyrosine-autophosphorylation site on c-KIT and PDGFR suggesting cellular efficacy is maintained (Figs 3B&C). Interestingly, titration of Dyr726 on PDGF-BB treated pediatric DMG lines HSJD-DIPG-007 (Fig 3D) and SU-DIPG-VI (Fig 3E) led to complete ablation of both phospho-AKT and phospho-ERK. There was no effect on phospho-Thr3 H3 levels suggesting that, similar to BRAF, Haspin is likely not engaged by Dyr726 in cells (Fig 3E). We further conducted colony formation assays on primary GBM102 cells with or without treatment with sub-micromolar concentrations of Dyr726, crenolanib, avapritinib, nilotinib, and imatinib. Dyr726 exhibited the most potent anti-glioma activity in a GBM102 colony formation assay (Fig 3F). Since PDFGRA has been reported to regulate mitophagy, we used the mitophagy reporter cell line ARPE-19 expressing the mito-QC reporter. As reported previously, Crenolanib induced significant mitolysosome punctae formation compared to DMSO-control (Fig 3G). Dyr726 mimicked the crenolanib phenotype with high accumulation of mitolysosome punctae suggesting potent upregulation of mitophagy (Fig 3G). This shows that Dyr726 is a highly potent inhibitor of PDGFRA.

**Figure 3:**
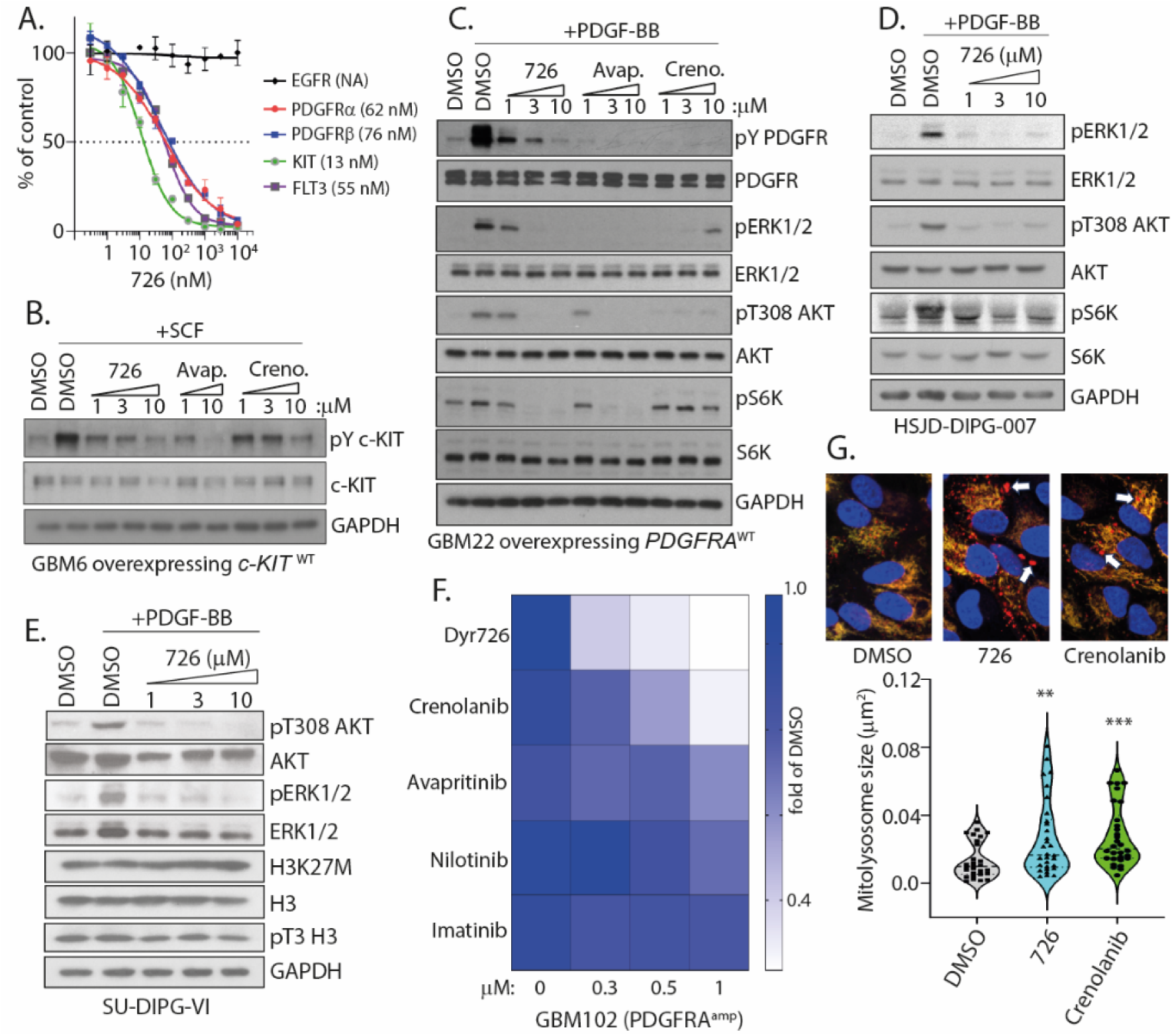
Dyr726 inhibits type III RTK signaling (A) EGFR, PDGFRα, PDGFRβ, KIT, FLT3 were assayed using the indicated concentrations of Dyr726. The IC_50_ graph was plotted using GraphPad Prism software. (B) GBM6 cells overexpressing c-KIT WT were serum starved for 2hr, then treated with SCF and indicated inhibitors. 2hr later cells were lysed and immunoblotting was performed using the indicated antibodies. (C) GBM22 cells overexpressing PDGFR WT were serum starved for 2hr, then treated with PDGF-BB and indicated inhibitors. 2hr later cells were lysed and immunoblotting was performed using the indicated antibodies. (D) HSJD-DIPG-007 cells were treated with PDGF-BB and the indicated concentration of Dyr726 or DMSO control for 2hr. Cells were then lysed and immunoblotting was performed using the indicated antibodies. (E) SU-DIPG-VI cells were treated with PDGF-BB and the indicated concentration of Dyr726 or DMSO control for 2hr. Cells were then lysed and immunoblotting was performed using the indicated antibodies. (F) GBM102 cells were treated with the indicated concentrations of either Dyr726, crenolanib, avapritinib, nilotinib, imatinib, or DMSO control for 7d. After 7d, cells were fixed with 100% EtOH and stained with crystal violet solution. 24hr later, the crystal violet solution was reconstituted using 10% acetic acid. Absorbance was measured at 590nm using the TECAN plate reader. Viability was measured as a fold change compared to DMSO treated cells. (G) ARPE-19 mitoQC cells were treated with 300nM Dyr726, Crenolanib, or DMSO for 24hr. Cells were then stained with Hoechst and mitolysosomes were imaged. Images were analyzed using the macro mito-QC Counter in FIJI. Mitolysosome size was quantified, ** P > 0.01 *** P < 0.001 (one-way ANOVA, mean +/- SD from n = 3 biological replicates).

### Dyr726 targets oncogenic gain-of-function mutants of PDGFRA and PI3Kα

As seen in Fig 1B and 2D, Dyr726 potently kills cancer cells which harbor PI3Kα and PDGFRA activating mutations. Since PI3Kα is one of the most mutated genes in cancer which is known to drive drug-resistance, we wanted to ascertain if Dyr726 could target cells harboring oncogenic mutations of PDGFRA and PI3Kα. We generated primary GBM22 cells stably expressing key activating mutations of PI3Kα and benchmarked the potency of Dyr726 with alpelisib and osimertinib (EGFR^MUT^ inhibitor). Compared to both alpelisib and osimertinib, Dyr726 exhibited enhanced potency in killing GBM22 cells with all PI3K activating mutants (Fig 4A). Alpelisib is FDA-approved currently for breast cancer, hence, we utilized immortalized mammary epithelial cells MCF10A parental or those harboring Crispr/Cas9 mediated knock-in of PI3Kα E545K and H1047R for benchmarking efficacy of Dyr726 against alpelisib. Titration of Dyr726 or alpelisib or lapatinib (EGFR/Her2 inhibitor) led to complete ablation of phospho-AKT, in all the MCF10A sub-clones (Fig 4B). Furthermore, Dyr726 potently targeted the cells and reduced colonies compared to alpelisib or lapatinib (Fig 4C&D). Next, we looked at PDGFRA activating mutants and generated GBM22 cells stably overexpressing PDGFRA D842V, Y288C, T674I, P345S, and V561D mutants. The mutants, especially D842V and V561D, are known to confer drug resistance to other type III RTK inhibitors like imatinib. Immunoblots confirm that avapritinib and Dyr726 both ablate phospho-AKT signal with T674I mutant being most resistant to both inhibitors (Fig 4E). However, Dyr726 is more potent than avapritinib at reducing the colonies of the mutants (Fig 4F). Interestingly, crenolanib was marginally more potent in reducing colonies than Dyr726 (Fig 4F).

**Figure 4:**
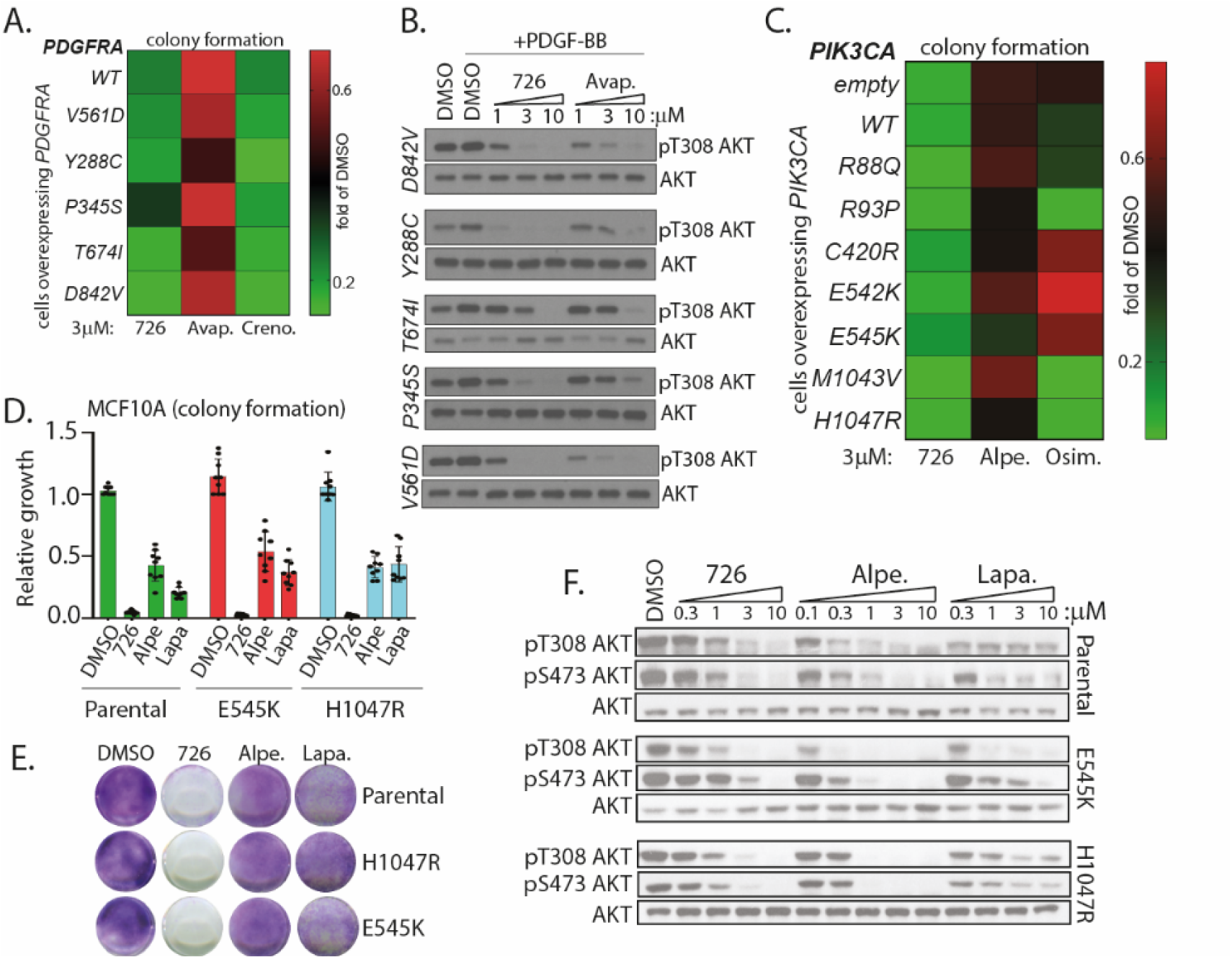
Dyr726 targets PDGFRA and PI3K activating mutations (A) GBM22 WT or mutant overexpressing PDGFRA cells were treated with either 3µM Dyr726, avapritinib, crenolanib, or DMSO control for 7d. After 7d, cells were fixed with 100% EtOH and stained with crystal violet solution. 24h later, the crystal violet solution was reconstituted using 10% acetic acid. Absorbance was measured at 590nm using the TECAN plate reader. Viability was measured as a fold change compared to DMSO treated cells. (B) GBM22 cells overexpressing PDGFRA WT, D842V, Y288C, T674I, P345S, V561 were serum starved for 2hr, then treated with PDGF-BB and inhibitor. 2h later cells were lysed and immunoblotting was performed using the indicated antibodies. (C) Indicated GBM22 WT or mutant cells were plated at 10,000 cells/well. 24hr later they were treated with 3µM of Dyr726, alpelisib, Osimertinib, or DMSO control. 1 week later, cells were fixed with 100% EtOH and stained using crystal violet solution. 24h later, the crystal violet solution was reconstituted using 10% acetic acid. Absorbance was measured at 590nm using the TECAN plate reader. Viability was measured as a fold change compared to DMSO treated cells. (D) MCF10A WT, H1047R, or E545K mutant cells were treated with either DMSO, or Dyr726 or alpelisib, or lapatinib. After 7d, cells were fixed with freshly made 4% paraformaldehyde (PFA). Cells were stained with crystal violet solution and then left to air dry overnight. Plates were scanned and then FIJI was used to analyze colonies. This was performed by cropping to an individual plate and converting to a binary image. The fill holes, watershed and analyze particles functions were then used to count colonies. (E) Representative images from (D) shown. (F) MCF10A WT, E545K, or H1047R mutant cells were treated with indicated concentrations of Dyr726, alpelisib, or lapatinib for 1hr. Cells were then lysed and immunoblotting was performed using the indicated antibodies.

### Dyr726 ablates WNT signaling through inhibition of DYRK/CLK kinases

Dyr726 is a very potent inhibitor of the DYRK and CLK family of CMGC kinases with biochemical IC_50_s of less than 100 nM for all isoforms except CLK3 (277 nM) (Fig 5A). Dyr726 does not inhibit GSK3β (Fig 5A). Indeed, Dyr726 treatment reduces 26S proteasome chymotryptic- and tryptic-like activity in cells by 20-30% suggesting DYRK2 engagement in cells (Fig 5B). Similarly, titration of Dyr726 ablates SF3BP1 phosphorylation on Thr474 suggesting DYRK1A/B engagement in cells (Fig 5C). Pan-inhibition of DYRK/CLK has been reported previously to drive WNT inhibition and hence we tested the ability of Dyr726 to inhibit the WNT signaling pathway using TopGFP expressing human colon epithelial cells (HCEC). Dyr726 titration markedly inhibited WNT signaling in either WNT3A activated cells or in cells treated with GSK3β inhibitor CHIR99021 (Fig 5D). Historically WNT-pathway inhibitors have failed in clinical trials for toxicity reasons, so we benchmarked Dyr726 WNT inhibition and HCEC reporter cell toxicity to previously reported WNT and DYRK/CLK inhibitors. Interestingly, Dyr726 was much more potent in inhibiting the WNT pathway compared to other previously reported WNT or DYRK/CLK inhibitors while being less toxic to the non-cancerous reporter HCEC cells. This suggests that Dyr726 is the most potent yet least toxic WNT pathway inhibitor in this table (Fig 5E). Intriguingly, abemaciclib was also found to be a potent WNT pathway inhibitor, an activity likely attributed to the DYRK/CLK inhibition as reported previously (Fig 5E). Next, we tested if Dyr726 reduced the transcription of WNT-targets and utilized a WNT qPCR array. We treated GBM22 cells with Dyr726 at 10 µM for 24h. Indeed, all WNT target gene transcripts were down (Fig 5F). WNT-target genes AXIN2, c-MYC, DVL2, BTRC, and LRP5 transcripts were reduced in both adult GSCs and in gliomagenic murine neural stem cells NPE (Fig 5G). We further observed that treatment with 10 μM Dyr726 over 24h led to downregulation of c-MYC protein levels in Myc-amplified GBM22 cells (Fig 5H). Overall, Dyr726 is a highly potent WNT pathway inhibitor through specific CMGC kinase inhibition and exhibits reduced toxicity compared to all other WNT inhibitors tested in the clinical trials.

**Figure 5:**
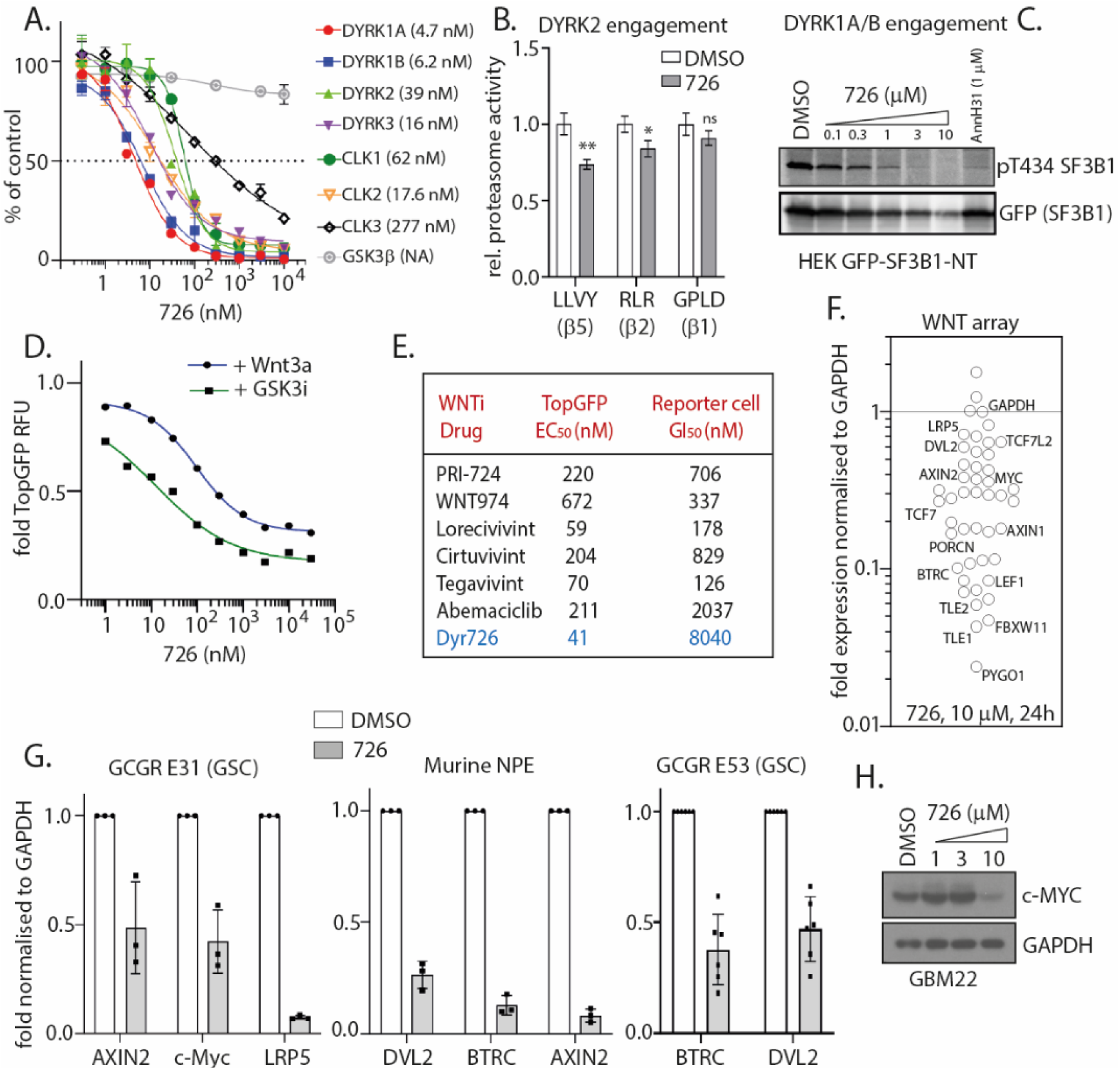
Dyr726 ablates the WNT pathway through DYRK/CLK inhibition. (A) DYRK1A, DYRK1B, DYRK2, DYRK3, CLK1, CLK2, CLK3, and GSK3β were assayed using the indicated concentrations of Dyr726. The IC50 graph was plotted using GraphPad Prism software. (B) Proteasome activity in total cell lysates from GBM22 cells with or without 10 μM 726 treatment for 2 h was measured with Suc-LLVY-AMC or Ac-RLR-AMC or Ac-GPLD-AMC. *P < 0.05, **P < 0.01 (compared to control treated, 2-way ANOVA, mean ± SD from n = 3 biological replicates). (C) HEK cells transiently expressing GFP-SF3B1-NT were treated with the indicated concentrations of 726 or 1 μM AnnH31 for 48 h. The phosphorylation state of SF3B1 was determined by immunoblotting with pThr434 antibody, and total amount of GFP-SF3B1 was utilized as loading control. (D) Comparison of Dyr726 treatment on TopGFP HCEC WNT reporter assay with or without either CHIR99021 (+GSK3i) or WNT3A treatment. The EC_50_ graphs were plotted as mean±SEM using GraphPad Prism software. (E) TopGFP WNT reporter assay was utilized to benchmark inhibitory activity of 726 against known WNT inhibitors that has undergone clinical trials. (F) Relative expression of WNT target genes were carried out in a WNT qPCR array in GBM22 cells upon 10 μM Dyr726 treatment for 24 h. (G) Relative expressions of indicated transcripts were carried out in the indicated cells upon 10 μM 726 treatment for 24 h. (H) GBM22 cells treated with or without indicated concentrations of Dyr726 for 18 h. Later cells were lysed and immunoblotting was performed using the indicated antibodies.

### Dyr726 kills WNT-dependent cancer organoids and extends survival of tumour mice *in vivo*

Since Dyr726 is a WNT-PI3Kα-pathway inhibitor, we wanted to test the efficacy of the molecule against a cancer model which would harbor both pathways for survival and proliferation. Hence, we acquired a library of 10 WNT-dependent colorectal cancer (CRC) patient-derived tumour organoids (PDOs) for testing compound efficacy *ex vivo*. These PDOs are epithelial adenocarcinomas with very high WNT pathway activity, that contain heterogeneous populations of cancer stem cells (often with PI3Kα GOF mutations), proliferative cells, and differentiated post-mitotic cells, including goblet, colonocyte, and endocrine cells. We tested Dyr726 across the PDO library with diverse driver mutation profiles and observed cytotoxic effects on most of the PDOs (Fig 6A&B). We observed a range of effects, with some PDOs being very sensitive to Dyr726 in the sub-300 nM range (588824), while others showed sensitivity to higher 3 μM doses (Fig 6A&B). Interestingly, one PDO (282377) was intrinsically resistant to Dyr726 even at a 3 μM dose (Fig 6A&B). This data highlights the effect of Dyr726 on *ex-vivo* tumour models that more closely capture tumour heterogeneity observed *in vivo*. Encouragingly on dosing in mice intraperitoneally at 25 mg/kg, Dyr726 exhibited excellent brain penetrance Kp (∼1) and Kp,uu (0.34) (Fig 6C). Physiologically relevant unbound brain concentration of Dyr726 was above the PI3Kα IC_50_ for > 24 h and above the WNT (EC_50_) and PDGFRA (IC_50_) for 8 and 4 h respectively (Fig 6C). As such dosing Dyr726 at 25 mg/kg was an appropriate dose to probe our multi-targeted hypothesis in rodent glioma models. To determine the efficacy of Dyr726 *in vivo*, an orthotopic allograft was generated using GL261 cells and mice were treated with either vehicle, Dyr726, or abemaciclib (Fig 6D). Compared to vehicle, Dyr726 treatment significantly improved and almost doubled overall survival of glioma-bearing mice. Abemaciclib was also effective at a similar concentration although Dyr726 treated mice had an overall improved survival (Fig 6D). Similar improvement in survival was observed in orthotopic GBM12 patient-derived xenograft mice wherein Dyr726 treated cohort exhibited improved overall survival compared to vehicle treated control upon 12 doses of 25 mg/Kg Dyr726 (Fig 6E).

**Figure 6:**
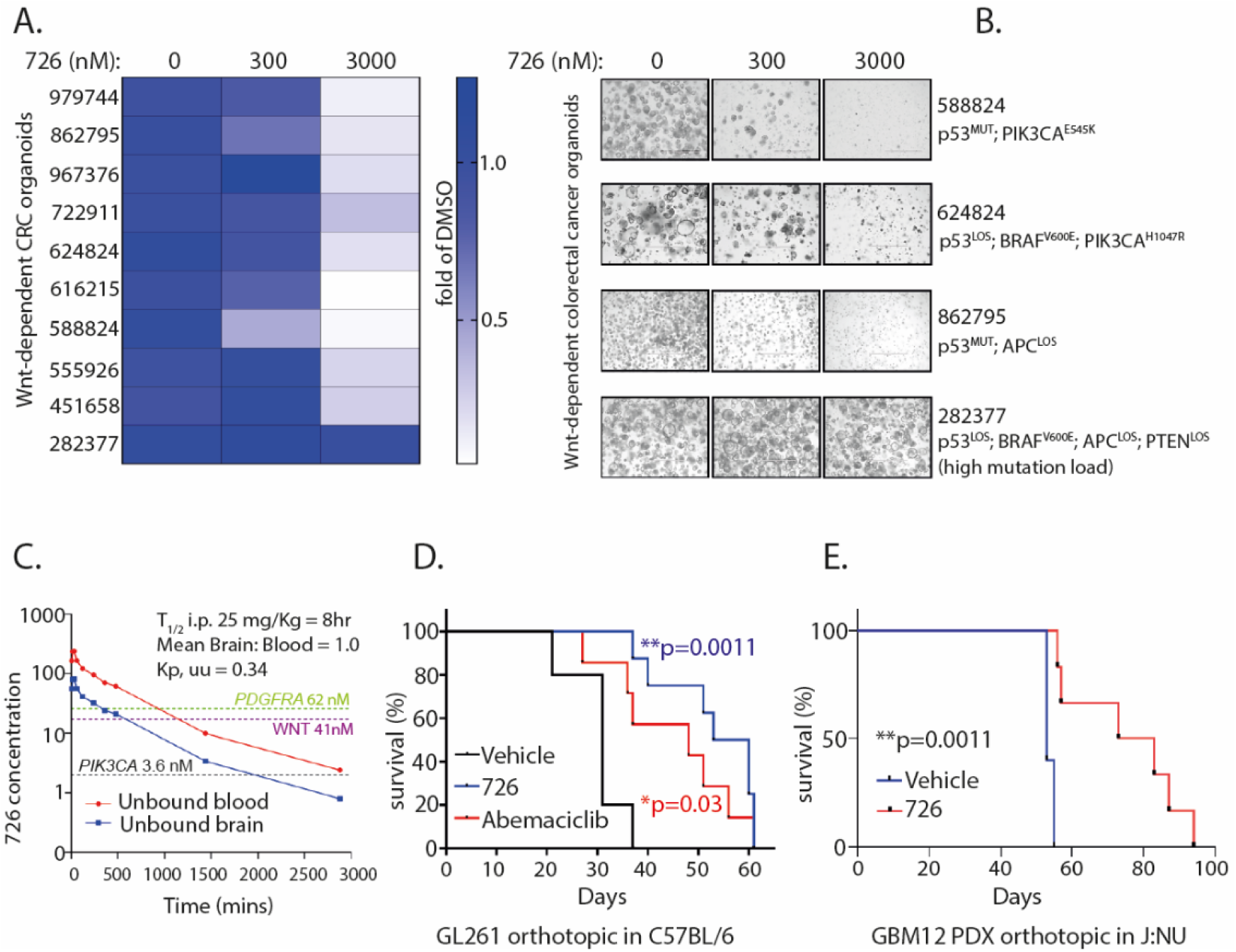
Dyr726 reduces *ex vivo* organoid and *in vivo* tumour volume (A) Indicated WNT-dependent patient-derived CRC organoids were treated with indicated concentrations of Dyr726 and cell viability were measured after specific time points using CellTiter-Glo® 3D reagent. Data was generated as fold of untreated form 3-6 biological replicates. (B) Representative images of specific organoids from (A) shown (C) Pharmacokinetic profile of Dyr726. Unbound brain exposure (ng/mg) with unbound blood exposure (ng/mL) are plotted at the indicated times (x-axis) following a single dose of 25 mg/kg (nL=L3 mice per time point). The biochemical IC_50_ for each of the key targets is overlayed with the PK plot. Also see Supplementary Fig S2. (D) 2,000 GL261 cells were surgically injected intracranially into C57BL/6 mice of either gender. 5d post-surgery, mice were randomized and treated with either vehicle (n=14), Dyr726 25mg/kg (n=8), or abemaciclib 25 mg/kg (n=8) 3x weekly. At designated endpoint of 10% weight-loss, mice were culled, and Kaplan-Meier curve was derived (P value derived from survival curve comparison using Mantel-Cox Log-rank test). (E) 12,000 GBM12 cells were surgically injected intracranially into J:NU mice of either gender. 14d post-surgery, mice were randomized and treated with either vehicle (n=5), or Dyr726 25 mg/kg (n=6) 3x weekly. At designated endpoint of 10% weight-loss, mice were culled, and Kaplan-Meier curve was derived (P value derived from survival curve comparison using Mantel-Cox Log-rank test).

## Discussion

The current work demonstrates a multidisciplinary approach in developing a first-in-class brain-penetrant PI3Kα, type III RTK, and WNT pathway inhibitor for therapeutic targeting of specific cancers of un-met need. The compound was discovered after extensive SAR development on our previously reported brain-penetrant DYRK1A inhibitor Dyr219 developed for targeting phospho-tau in Alzheimer’s disease^21–24^. Dyr726 exhibits good brain penetrance and efficacy. Compared to most of the WNT pathway inhibitors tested in the clinical trials, Dyr726 shows improved efficacy and engagement while maintaining a therapeutic window of over 200-fold as judged by its WNT activity versus toxicity in HCEC cells. Abemaciclib, an FDA-approved CDK4/6 drug, is a relatively modest WNT pathway inhibitor driven by its DYRK/CLK inhibitory activity. Note abemaciclib’s WNT inhibitory activity is somewhat controversial as groups have reported WNT pathway activation through off-target GSK3 inhibition^25^. Dyr726 is a potent WNT pathway inhibitor through DYRK/CLK engagement, with no off-target activity on GSK3β. Furthermore, Dyr726 has a much-improved PI3Kα engagement profile with potent ablation of PI3K-mTOR pathway across primary glioma, DMG, breast cancer cells either harboring parental or gain-of-function mutations of PI3Kα. PI3Kα H1047R mutation has been reported to drive lapatinib resistance which is partly observed in our hands^26^. However, Dyr726 ablated all cancerous cells regardless of mutation status. This is thought largely due to the desirable multi-targeted profile and not inherent toxicity as mice treated with over 12 doses of Dyr726 at 25 mg/kg do not exhibit weight loss or hyperglycemia (Supplementary Fig S2) which are often reported for PI3K and WNT pathway inhibitors^12, 27^. It is also important to note that maximal tolerated dose of Dyr726 was greater than 50 mg/Kg and is AMES negative thus has no mutagenic properties either. The ability of Dyr726 to target type III RTKs and especially PDGFRA suggests that the compound could target DMG thus expanding the repertoire to beyond adult GBM or WNT-dependent CRC. This is evident with Dyr726 targeting 3D cultures of DMG lines. While Dyr726 does not affect the epigenetic driver of H3.3 mutation, its ablation of key oncogenic drivers could provide an alternative treatment approach for DMG which currently has no chemotherapeutic options available. It is important to note that in all EGFR driven tumour models, Dyr726 ablates mTOR signaling but does not exhibit any inhibitory effect in the MAPK pathway as seen by a modest increase in pERK. Since EGFR is not inhibited by Dyr726, there is a chance that EGFR-MAPK could drive a potential resistance mechanism against Dyr726. The only evidence of intrinsic resistance we observed was in a WNT-dependent CRC organoid which harbored over 10 compounding driver mutations. Hence, we will focus more on intrinsic and adaptive resistance pathways to Dyr726 as we progress further down the translational pipeline toward a clinical trial. Importantly, PDGFRA driven DMG exhibits a complete ablation of both mTOR and MAPK pathways which further strengthens the hypothesis that Dyr726 could be potentially beneficial for DMG. Regardless, Dyr726 doubled the overall survival of orthotopic glioma allo- and xenograft mice with no major toxic outcomes. It could be argued that Dyr726 may be more beneficial than clinical drug abemaciclib in targeting orthotopic glioma.

Taken together, the remarkable kinome specificity, in conjunction with its select multitargeted profile, drives the efficacy and safety profile of Dyr726 which not only works well in combination with glioma standard-of-care *in vitro* but also exhibits improved cell-kill over key clinicals which have undergone or are currently undergoing brain cancer clinical trials. Overall, the drug discovery campaign encompassing Dyr762 and close analogs shows promise as the only known PI3K-mTOR, WNT-pathway and type III RTK inhibitor with good brain penetrance and no visible toxicity while reducing tumour burden. Efficacious tumour reduction *ex vivo* and *in vivo* by Dyr726 provides a testament to how a controlled and benchmarked multi-targeted approach to drug discovery could improve our understanding of cancer biology, mechanisms of resistance and how to drive brain-penetrant congener exploration to potentially avoid toxicity going forward.

## Acknowledgements

The authors thank the Kinase-Screen team at International Centre for Protein Kinase Profiling, Biological services team at the Animal Resource Unit, and the Fingerprints Proteomics team at the University of Dundee, UK.

---

**Supplementary Fig S1.**
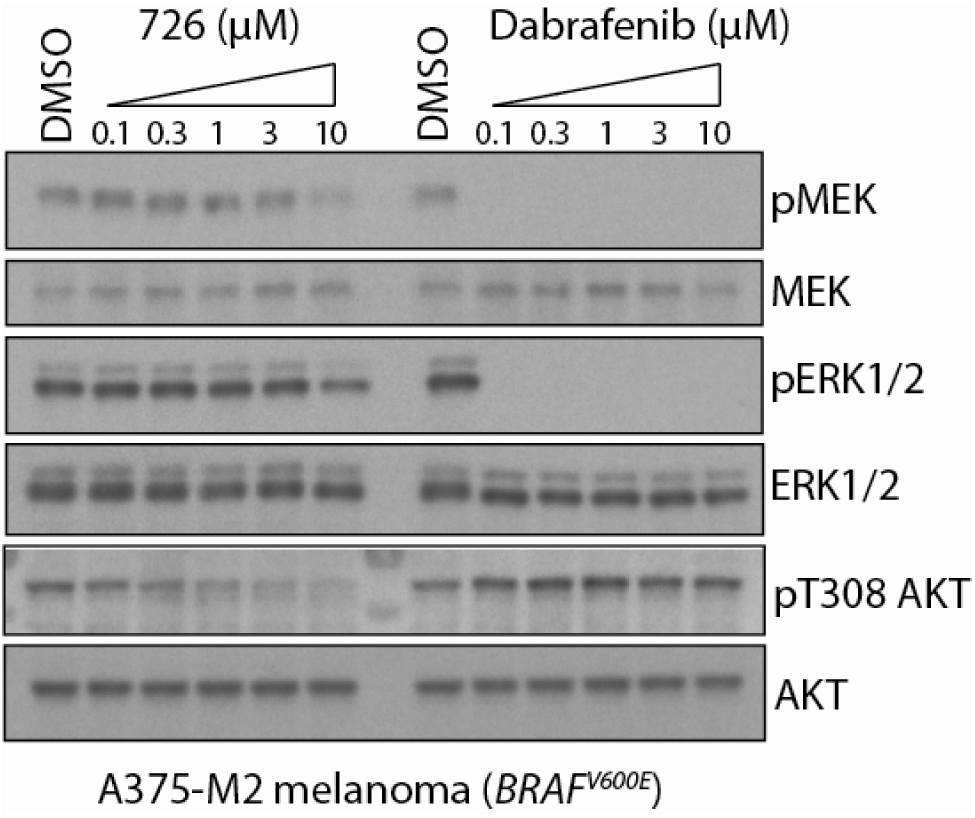
Cellular BRAF engagement by Dyr726. A375-M2 melanoma cells harbouring endogenous BRAF V600E mutation were treated with the indicated inhibitors. 2hr later cells were lysed and immunoblotting was performed using the indicated antibodies.

**Supplementary Fig S2.**
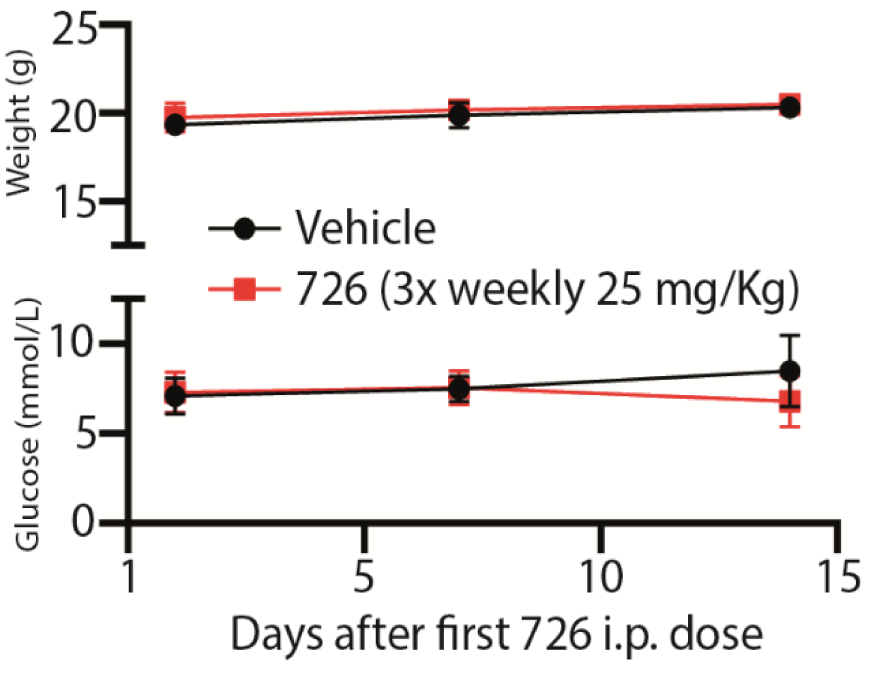
Dyr726 treatment does not affect the overall body weight or blood glucose levels in mice over 2 weeks of 3x weekly 25 mg/Kg Dyr726 treatment.

## Materials and Methods

### Cell culture growth conditions

Mammalian cells were all grown in a humidified incubator with 5% CO_2_ at 37 °C. Cells were dissociated with Accutase®Lsolution (GCGR cell lines), TrypLE™ Express Enzyme (GBM Neurospheres) or Trypsin-EDTA (0.05%) once flask reached 80% confluency. NPE and GCGR cell lines (E31, E53) were grown in DMEM/HAMS-F12 supplemented with 1.5% (v/v) Glucose, 1% (v/v) MEM Non-Essential Amino Acids Solution, 1% penicillin-streptomycin, 0.16% BSA (v/v), 0.2% β-mercaptoethanol (v/v), B27 supplement, N2 supplement, 10 ng/ml human EGF, 10ng/ml human FGF, 2ug/ml Laminin and 1% penicillin-streptomycin^28^. GBM cell lines (GCE40, GIN40, GCE41, GBM22, GBM6, GBM10, GBM44, GBM64, GIN31, GCE31) were cultured in DMEM supplemented with 10% Fetal Bovine Serum (FBS), 1% penicillin-streptomycin, 10ug/ml insulin, 20ng/ml human EGF. Human Colonic Epithelial Cells (HCEC) were cultured using 1x DMEM supplemented with 1% penicillin/streptomycin, 1% Glutamax, and 10% FBS. A375-M2 melanoma cells and BT20 cells were cultured in DMEM and EMEM respectively, supplemented with 10% FBS, and 1% penicillin-streptomycin. G7 cells were maintained on Matrigel coated corning plastic in Advanced DMEM, hEGF, hFGF, B27 supplement (1%), N-2 supplement (0.5%), Glutamine 2mM, heparin (5ug/ml). MCF10A WT, MCF10A PI3K E545K, and MCF10A PI3K H1047R cells were cultured in DMEM/F-12 medium supplemented with 5% horse serum, 20 ng/ml hEGF, 0.5 μg/ml hydrocortisone, 100 ng/ml cholera toxin, 10 μg/ml insulin and 1% penicillin and streptomycin. HSJD-DIPG-007 and SU-DIPG-VI neurospheres were cultured in base media containing equal parts Neurobasal A and DMEM/F12 supplemented with 10mM HEPES buffer, 1mM MEM sodium pyruvate solution, 1X MEM Non-Essential Amino Acids, 1X GlutaMAX-I Supplement, 1X Antibiotic-Antimycotic, and supplemented with fresh B27 without vitamin A, 2 μg/mL heparin, 20 ng/mL human-EGF, 20 ng/mL human-bFGF, 10 ng/mL PDGF-AA, and 10 ng/mL PDGF-BB. GBM neurospheres (GBM120, GBM12, GBM76, GBM192, GBM215) were cultured in Knockout DMEM/F-12 Basal Media supplemented with StemPro NSC SFM supplement, 20ng/ml hEGF, 20ng/ml hFGF, and 1% penicillin-streptomycin. ARPE-19 mito-QC reporter cells were cultured in 1:1 DMEM:F-12 media supplemented with 10 % (v/v) FBS, 2 mM L-glutamine, 100 U/ml penicillin and 0.1 mg/ml streptomycin. Human colonic epithelial cells (HCECs) were cultured in DMEM treated with 10% FBS, 1% Glutamax and 1% penicillin/streptomycin. We obtained patient-derived organoids from the National Cancer Institute Patient-Derived Model Repository (PDMR; NCI-Frederick, Frederick National Laboratory for Cancer Research, Frederick, MD, USA). The PDOs were expanded, sub-cultured, and cryopreserved following the NCIs standard operating procedures (SOPs) 40102, 40103, 40104 and the media requirements from SOP 30101.

### Drug reconstitution

For in vitro experiments, all drugs were dissolved in DMSO at a stock concentration of 10 mM. For *in vivo* treatments, Dyr726 and abemaciclib were formulated as a solution at a concentration of 2.5 mg/mL, in a mixture of 5% DMSO (dimethyl sulfoxide), 47.5% PEG400 (Polyethylene glycol 400), and 47.5% saline (0.9% w/v NaCl). To produce this formulation, Dyr726 or abemaciclib was weighed using a calibrated analytical balance and transferred to a squat glass vial with a magnetic stir bar. Dimethyl sulfoxide was added, and the resulting mixture was stirred vigorously until a fine, homogenous suspension was obtained. Polyethylene glycol 400 was then added and stirring continued for 10-15 minutes. Once a complete solution had been obtained, the stir speed was reduced to a maximum of 300rpm. The required volume of saline was added dropwise over the course of 5-10 minutes. The resulting solution was transferred into aliquots for dosing, and stored at 2-4°C.

### *In vivo* DMPK studies – Pharmacokinetics and Brain Penetration

Mouse pharmacokinetics and drug brain penetrance experiments were carried out under the authority of Project License Number PP5016780 granted by the UK Home Office under the Animals (Scientific Procedures) Act 1986. Prior to submission to the Home Office, applications for project licenses were approved (approval number WEC2019-08) by the University Welfare and Ethical Use of Animals Committee, acting in its capacity as an Animal Welfare and Ethical Review Body as required under the Act.

Nine female mice of the C57BL/6J strain were obtained from Charles River Laboratories, UK. Animals were maintained under a 12-hour light / 12-hour dark cycle, in Thoren mouse caging containing Chips7D bedding (Dates and Group, UK), with ad libitum access to food (RM1; Special Diet Services, UK) and water. Temperature and relative humidity were maintained between 20-24°C and 45-65% respectively. All animals receive a minimum of 14 days acclimatization prior to the start of the study. Dose formulations were prepared on the day of dosing.

For the pharmacokinetic study, the formulation was dosed to three mice via intraperitoneal injection at a dose level of 25 mg/kg (10mL/kg). Animals were observed regularly after dose administration. Serial blood samples (10 µL per sample) were collected from the lateral tail vein prior to dosing, then at 5, 15 and 30 min, 1, 2, 4, 6, 8, 24 and 48 hours after the administration of Dyr726. Each blood sample was diluted into 90 µL of Milli-Q ultrapure water and stored at 20°C prior to bioanalysis. The concentration of Dyr726 in whole blood was determined via UPLC-MS/MS.

For the brain penetration study, the formulation was dosed to six mice via intraperitoneal injection at a dose level of 25 mg/kg (10mL/kg). Animals were observed regularly after dose administration. At 30min, and 4.5 hours after the administration of Dyr726, 3 animals per time point were placed under terminal anesthesia with isoflurane. Blood was collected via cardiac puncture and the brain removed. Two aliquots of 10 µL blood were diluted into 90 µL of Milli-Q ultrapure water. All samples were stored at 20°C prior to bioanalysis. The concentration of Dyr726 in whole blood and brain tissue was determined via UPLC-MS/MS.

### Orthotopic animal experiments

All mouse orthotopic tumour experiments were carried out under the authority of Project License Number PP2096524 granted by the UK Home Office under the Animals (Scientific Procedures) Act 1986. Prior to submission to the Home Office, applications for project licenses were approved (approval number WEC2022-06) by the University Welfare and Ethical Use of Animals Committee, acting in its capacity as an Animal Welfare and Ethical Review Body as required under the Act.

Intracranial transplantation of cells was performed using a stereotaxic frame to inject the cells resuspended in 5μl of distilled Phosphate buffered saline (PBS) into mice after administration of isoflurane general anesthesia and analgesic buprenorphine (Vetergesic) 0.1mg/kg subcutaneously. 2,000 GL261 cells were intracranial transplanted into C57BL/6. One-week post-operation, mice were randomized into three groups of either vehicle (n=14), DYR726 25 mg/kg (n=8), or abemaciclib 25 mg/kg (n=8) 3x weekly. 12,000 GBM12 neurospheres were intracranial transplanted into 9-week-old of 12 athymic nude mice (obtained from Charles River, UK) striatum. One-week post-operation, mice were randomized into two groups n=6 to begin treatment of either vehicle or DYR726 at 25 mg/kg via intraperitoneal injections. End point was determined by >10% body weight loss or a body condition score of 2 on the standard 5-point scale. One mouse from the vehicle cohort developed an ectopic tumour outside the skull and was removed from the study.

### DNA constructs and Cell Transfection

pHAGE-PDGFRA was a gift from Gordon Mills & Kenneth Scott^29^ and was used as the parental vector for mutagenesis. Mutant plasmids were generated by point mutation via site-mutagenesis using the Q5 Side-Directed mutagenesis Kit (New England Biolabs) according to the manufacturer’s instructions. Mutations were confirmed by Sanger sequencing. Mutation primers are listed in Table X. The GeneJET Plasmid Miniprep Kit was used to extract the plasmids from transformed DH5α bacterial competent cells.

Transient transfection was carried out using Lipofectamine 3000 (Life Technologies) or Polyethylenimine “Max” (PEI MAX) as recommended by the manufacturer. Stable overexpression of TopGFP reporter/H2BmCherry, PDGFRA and PI3K WT and mutant constructs were carried out by lentivirus transfection followed by GFP selection. For lentivirus production, HEK293T Phoenix cells were transfected at 70-80% confluency using Lipofectamine 3000 and psPAX2 and pMD2.G packaging vectors. Medium was changed 4 hours after transfection and supernatant was collected after 72 hr. Viral media collected and filtered (45μm) and mixed with 8 µg/ml Polybrene before being added to recipient cells and left to incubated for 48h.

### Cell viability assay

Cell viability was measured as stated previously^30, 31^. Briefly, equal number of cells (5,000 cells/well) were seeded per well in 96 well plates. Cell viability assays were carried out with 72 hr treatment of indicated drugs or DMSO control using CellTiter 96® AQueous Non-Radioactive Cell proliferation assay, adhering to manufacturer instructions. Absorbance was measured using a Tecan multi-well plate reader and data was represented as % viability compared to DMSO treated control.

For cell viability experiments using 3-dimensional organoid experiments, organoids were harvested using Dispase II according to SOP 40103. Organoids were dissociated to single cells using 0.25% Trypsin/EDTA, and the cells were counted. Cells were plated in 10 µl Basement Membrane Extract domes (Cultrex UltiMatrix) in 48-well plates at a density of 5,000 to 10,000 cells/dome. Two to three domes were plated for each treatment condition. Cells were incubated overnight in growth medium plus Y-27632 dihydrochloride and CHIR99021 at 10 µM and 2.5 µM, respectively. The next day, the plating medium was removed and replaced with growth media containing DMSO or DYR726 compounds in concentrations of 30, 300, or 3000 nM. Hydroxyurea at 2 mM was used as a control that would strongly promote organoid death. Organoid media was changed every 3 to 4 days. Prior to performing the CellTiter-Glo® 3D Viability Assay, organoid domes were imaged on an EVOS FL Auto microscope with Pearlscope 64 software using a 4X objective. For the viability assay, the media was shaken off the plates, and 100 µl of 2X Passive Lysis Buffer (diluted from 5X) was added to each well. The domes were loosened using a pipet tip and the plate was shaken at 300 rpm for 30’ at room temperature. After 30 minutes, the loosened domes were transferred to an opaque, white, 96-well assay plate. 100 µl of the CellTiter-Glo® 3D reagent, diluted 1:10 into 2X Passive Lysis buffer, was added to each well, and the plate was shaken again for 30 minutes, after which luminescence was read on a plate reader.

### Spheroid invasion assay

3D spheroid invasion assay was carried out as stated previously^32, 33^. Briefly, 2,000 U87.MG or GBM102 cells were seeded per well in a low attachment u-bottom 96 well plate and spheroids were allowed to form over four days. These spheroids were embedded in Geltrex which is a basement membrane extract that contains laminin, collagen IV, entactin, and heparin sulphate proteoglycans. The embedded spheroids were supplemented with complete media at top with or without drugs. The photomicrographs of each well containing spheroid were taken at x100 resolution starting from day 0 (day of embedding the spheroid in matrix) through day 4 or day 6. The relative invasion was calculated using Image J software.

### Neurosphere formed and formation assays

Neurosphere targeting assays were carried out as stated previously^34^. Neurospheres were plated at 5,000-10,000 cells/well in a 96 well plate. For formation assays, on day 0, cells were either treated with DMSO or the respective drugs. For formed assays, cells were kept in the incubator for 7 days to allow the neurospheres to form and either treated with DMSO or the respective drugs. Images were taken on day 0, 7, and 14 on the FLoid™ Cell Imaging Station. Neurosphere diameter was measured using FIJI.

### Colony formation assay

MCF10A cells were plated into 15 cm^2^ plates at 250 cells/ml^35^. Cells were then treated with either DMSO or the respective drugs. After 7 days, cells were fixed with 4% paraformaldehyde (PFA). Cells were stained with 0.1% Crystal Violet solution and then left to air dry overnight. The cells were dried, and images were captured using a microscope (Olympus, FV500-IX71, Japan) to count the number of colonies containingL>L50 cells. GBM22, GBM102, PI3KCA and PDGFR WT and mutant cells were plated into 12 well plates at 10,000 cells/ml. Cells were then treated with either DMSO, or the respective drugs. After 7 days, cells were fixed with 100% Ethanol and stained with 0.1% Crystal Violet solution. 24h later, the crystal violet solution was reconstituted using 10% acetic acid. Absorbance was measured at 580 nm using the TECAN plate reader. Viability was measured as a fold change compared to DMSO treated cells.

### *In vitro* Radiation experiments

The radio-sensitizing effects of Dyr726 were measured by clonogenic survival in the patient-derived glioma cells G7 (Colin Watts lab Cambridge). Cells were plated onto Matrigel coated 6 well plates (250 cells/well) and incubated overnight prior to treatment with DYR726. Cells were pre incubated with either DMSO or the respective drug for 1 hour prior to irradiation in an Xstrahl RS225 cabinet (Suwanee, GA, USA). 14 days post irradiation the colonies were fixed with methanol and stained with crystal violet solution [1:40 dil in PBS (Sigma HT90132)]. The plates were then scanned on an Oxford optromix gelcount (Adderbury, UK) and manually counted using ImageJ software (NIH, Bethesda, USA)

### IC_50_ determination and kinase specificity profiling

IC_50_ determination was carried out at The International Centre for Protein Kinase Profiling. (http://www.kinase-screen.mrc.ac.uk/). PI3Kα, PI3Kβ and PI3Kγ (diluted in 12.5mM Glycine-NaOH (pH 8.5), 50mM KCl, 2.5mM MgCl2, 1mM DTT, 0.05% CHAPS) was assayed in total volume of 20ul containing 12.5mM Glycine-NaOH (pH 8.5), 50mM KCl, 2.5mM MgCl2, 1mM DTT, 0.05% CHAPS, 0.01mM ATP and 0.05mM diC8 PIP2. The enzyme was assayed for 60 or 75 min after which 20ul of ADP-Glo reagent was added. After a further incubation of 40 min, 40ul of Kinase Detection Buffer was added. The assays were incubated for 40 min and then read on PerkinElmer Envision for 1sec/well.

CLK1, CLK2, CLK3 (5-20mU diluted in 50 mM Tris pH 7.5, 0.1 mM EGTA, 1 mg/ml BSA) was assayed against RNRYRDVSPFDHSR peptide in a final volume of 25.5 µl containing 50mM Tris pH 7.5, 0.3mM peptide, 10mM DTT, 10 mM magnesium acetate and 0.005 mM [33P-γ-ATP] (50-1000 cpm/pmole) and incubated for 30 min at room temperature.

DYRK1A, DYRK1B, DYRK2, DYRK3 (5-20 mU of diluted in 50 mM Tris pH7.5, 0.1 mM EGTA) was assayed against Woodtide (KKISGRLSPIMTEQ) in a final volume of 25.5µl containing 50 mM Tris pH 7.5, 0.1 mM EGTA, 350 µM substrate peptide, 10 mM Magnesium acetate and 0.05 or 0.005 mM [33P-γ-ATP](50-1000 cpm/pmole) and incubated for 30 min at room temperature^17^.

GSK3β (5–20 mU diluted in 20 mM MOPS pH 7.5, 1 mM EDTA, 0.01% Brij35, 5% glycerol, 0.1% β-mercaptoethanol, 1 mg/ml BSA) is assayed against Phospho-GS2 peptide (YRRAAVPPSPSLSRHSSPHQS(PO4)EDEEE) in a final volume of 25.5 µl containing 8 mM MOPS pH 7.0, 0.2 mM EDTA, 20 µM Phospho GS2 peptide, 10 mM magnesium acetate and 0.005 mM [33P-γATP] (50-1000 cpm/pmole) and incubated for 30 min at room temperature^17^. PDGFRα or PDGFRβ (5-20mU diluted in 50 mM Tris pH 7.5, 0.1 mM EGTA, 1 mg/ml BSA, 0.1% β-mercaptoethanol) was assayed against KKKKEEIYFFFG peptide in a final volume of 25.5 µl containing 50mM Tris pH 7.5, 0.1mM EDTA, 10mM DTT, 5mM MnCl_2_ 0.3mM substrate, 10 mM magnesium acetate and 0.005 mM [33P-γ-ATP] (50-1000 cpm/pmole) and incubated for 30 min at room temperature.

FLT3 (5-20 mU diluted in 50 mM Tris pH 7.5, 0.05% β-mercaptoethanol, 1 mg/ml BSA) is assayed against Catchtide (RRHYYYDTHTNTYYLRTFGHNTRR) in a final volume of 25.5 µl containing 50 mM Tris pH 7.5, 0.05% β-mercaptoethanol, 300 µM substrate peptide, 10mM magnesium acetate and 0.05 mM [33P-g-ATP] (50-1000 cpm/pmole) and incubated for 30 min at room temperature.

c-KIT (5-20 mU diluted in 50 mM Tris pH 7.5, 0.05% β-mercaptoethanol, 1 mg/ml BSA) was assayed against GGMEDIYFEFMGGKKK peptide in a final volume of 25.5 µl containing 50 mM Tris pH 7.5, 0.05% β-mercaptoethanol, 300 µM substrate peptide, 10mM magnesium acetate and 0.05 mM [33P-g-ATP] (50-1000 cpm/pmole) and incubated for 30 min at room temperature.

EGFR (5-20mU diluted in 50 mM Tris pH 7.5, 0.1 mM EGTA, 1 mg/ml BSA) is assayed against a substrate peptide (Poly Glut Tyr) in a final volume of 25.5 µl containing 50mM Tris pH 7.5, 1mg/ml substrate peptide, 10mM DTT, 2mM MnCl_2_, 10 mM magnesium acetate and 0.02 mM [33P-g-ATP] (50-1000 cpm/pmole) and incubated for 30 min at room temperature. CLK, DYRK, PDGFR, FLT3, c-KIT, and EGFR assays were stopped by addition of 5 µl of 0.5 M (3%) orthophosphoric acid and then harvested onto P81 Unifilter plates with a wash buffer of 50 mM orthophosphoric acid.

Kinase inhibitor specificity profiling assays were carried out at The International Centre for Protein Kinase Profiling and Eurofins (Supplementary Table 1). Dyr726 kinase specificity was determined against a panel of 156 protein and lipid kinases as described previously. The assay mixes and 33P-γATP were added by Multidrop 384 (Thermo). Results are presented as a percentage of kinase activity in DMSO control reactions. Protein kinases were assayed in vitro with 1 µM final concentration of Dyr726 and the results are presented as an average of triplicate reactions ± SD in the form of a waterfall plot.

### Western blot Analysis

Cells were lysed in SDS lysis buffer (250 mM Tris HCL pH 6.8, 2% (v/v) β-mercapto-ethanol, 2% SDS (w/v), 30% (v/v) Glycerol and 0.1% (w/v) bromophenol blue). Equal amounts of whole-cell lysates were subjected to gel electrophoresis using 10-12% SDS-PAGE (Tris-Glycine) and transferred to nitrocellulose membranes. After blocking in 5% Bovine serum albumin (BSA) or non-fat dry milk, membranes were incubated with the primary antibodies at 1:1000 or 1:200 for pT434 SF3B1. Following three washes in TBS-T, the blots were incubated with horseradish peroxidase-conjugated secondary antibody and the signals visualized by enhanced chemiluminescence system according to manufacturer’s instructions. Antibody details are described in the Materials table.

### Cell-based assay of DYRK1A/B inhibitory activity

The cellular efficacy of Dyr726 was assessed by measuring the phosphorylation of GFP-SF3B1-NT by endogenous DYRK1A/B, as described previously^36^. In brief, HEK tsA201 cells were transiently transfected with 500 ng/well GFP-SF3B1-NT in 6-well plates^37^. One hour after transfection, the cells were treated with Dyr726 or control compounds for 48 h. Subsequently, cell lysis was performed under denaturing conditions (100 µL/well 20 mM Tris-HCl pH 7.4, 1% SDS at 100°C). Phosphorylation of SF3B1 on Thr434 was detected by immunoblotting with a custom-made phospho-specific antibody ^37^. Relative phosphorylation levels of SF3B1 were determined by calculating the ratio of the pT434 signal to GFP immunoreactivity.

### Quantitative Tandem-Mass-Tag 6-plex Phosphoproteomics

#### Sample processing – Total proteome

GBM6, GBM22, and GIN28 cells were treated with either DMSO or 10 μM Dyr726 for 30 mins. Cells were lysed and samples were processed using S-Trap mini columns (Protifi) where proteins were digested with 16.7µg of trypsin per sample overnight and then a second digest at the same concentration for a further 6 hours. A fluorometric peptide quantification assay (Thermo Fisher) was used to quantify peptides prior to TMT labelling and 250µg of peptides per sample were taken for TMT labelling with a 6plex TMT kit (Thermo). Samples were subsequently desalted using Empore C18 cartridges and pooled into one sample. High pH reverse phase fractionation was performed on the pooled TMT sample resulting in 20 concatenated fractions ready for total proteome analysis.

<u>Mass Spectrometry Analysis – Total proteome:</u> Total Proteome fractions were run on a Q-Exactive HF (Thermo Scientific) instrument coupled to a Dionex Ultimate 3000 HPLC System (Thermo Scientific). A 5-35%B gradient compromising of eluent A (0.1% formic acid) and eluent B (80% acentonitrile/0.1% formic acid) was used to run a 120-minute gradient for fractions 1 to 10 and a 7-35%B gradient was used for fractions 11 to 20 with the same eluents and for the same length of time. Each fraction was loaded at 10 μL/min onto a trap column (100 μm × 2 cm, PepMap nanoViper C18 column, 5 μm, 100 Å, Thermo Scientific) equilibrated in 0.1% TFA. The trap column was washed for 5 min at the same flow rate with 0.1% TFA and then switched in-line with a Thermo Scientific, resolving C18 column (75 μm × 50 cm, PepMap RSLC C18 column, 2 μm, 100 Å) for the gradient. For the Full MS scan (MS1) data was collected in profile mode a resolution of 120,000, AGC target of 3E+06, maximum IT of 50ms and a mass range of 335-1600m/z was used with the top 15 most intense peaks in each MS1 scan were taken for MS2 analysis. The MS2 analysis collected data in centroid mode and used a resolution of 60,000, AGC target of 1E+05, maximum IT of 200ms and a fixed first mass of 100m/z. Spectra were fragmented using higher-energy C-trap dissociation (HCD).

#### Sample processing – Phosphopeptides

Total proteome fractions were pooled into 8 fractions for phosphopeptide enrichment. The phosphopeptide enrichment was carried out using Zr-IMAC beads using a 1:4 ratio.

<u>Mass Spectrometry Analysis – Phosphopeptides:</u> Phosphopeptide samples were run on an Orbitrap Fusion (Thermo Scientific) instrument coupled to a Dionex Ultimate 3000 HPLC System (Thermo Scientific). A 5-35%B gradient compromising of eluent A (0.1% formic acid) and eluent B (80% acetonitrile/0.1% formic acid) was used to a 130-minute gradient for fractions 1 to 8. For the Full MS scan (MS1) a resolution of 120,000, AGC standard target, maximum IT of 50ms and a mass range of 380-1500 m/z to acquire data in profile mode. Following the MS1 scan MS2 analysis was carried out in centroid mode using collision-induced dissociation (CID) with the CID energy set to 35%, activation time of 10ms, AGC standard target and a maximum IT of 60ms. Finally, MS3 analysis was collected using the following parameters; number of SPS precursor was set to 10, MS isolation window 0.7m/z, MS2 isolation window was set to 3m/z, activation type HCD and HCD collision energy of 65%, resolution of 60,000, scan range 100-500m/z, AGC target 200%, maximum injection time 105msand data was acquired in profile mode.

#### Data Analysis

TMT 6plex analysis was performed in MaxQuant version 1.6.6.0 using the RAW files generated with data manipulation in Microsoft Excel Office 365 to normalize the total proteome and phospho-peptide data as well as provide comparisons between samples. Volcano plot was generated using Graphpad Prism 10.

### Mitophagy

ARPE1-mitoQC cells were a kind gift from Prof Ian Ganley, University of Dundee^38^. Cells were plated into 18 well μ-Slides and treated with DMSO or the respective drugs. 24 hours later, cells were fixed with 4% PFA for 10min at 4*°C.* Cells were stained with Hoechst dye (Sigma #14533) at 200nM for 15 minutes at room temperature in the dark and then left in PBS + 0.2% Sodium Azide until mounting in Vectaschield non-setting medium. Cells were imaged with the Zeiss LSM 880 Airyscan microscope using a Plan Apochromat ×63 objective (NA 1.4) objective with a zoom of 1.0 and optical section thickness of 0.8µm (image size 1912 × 1912 pixels, pixel size 0.071µm). Images were analyzed using the macro mito-QC Counter in FIJI as described here.

### Proteasome activity assay

Proteasome activity assays were analyzed directly on cell lysates as described previously^31^. Briefly, cells were treated with or without indicated concentration of Dyr726 prior to lysis. Peptidase activities of purified proteasome or proteasomes in whole cell or tumour lysates were assayed using fluorogenic peptide substrates (Enzo Life Sciences) with phosphatase inhibitors present in the lysis and assay buffers. The measured activity was normalized either against total protein concentration of cell lysates. Fluorescence signals were quantified using a Tecan plate reader.

### Wnt reporter assay (Top flash GFP assay)

HCECs cells expressing the TopGFP reporter/H2BmCherry were seeded at 2,000 cells per well in 384-well black SCREENSTAR imaging microplates and allowed to adhere overnight. The following day, cells were stimulated to induce the Wnt pathway using 10μM CHIR99021 or 8 nM WNT3A. Simultaneously, DYR726 compound was delivered in a dose-response using a Tecan d300e digital dispenser ranging from 0μM to 30μM concentrations. Cells were incubated for 24 hours before fixing for 30 minutes with 4% paraformaldehyde/sucrose solution. Cells were permeabilized with 0.1% triton-x in PBS for 10 minutes and stained for DAPI for 30 minutes. Plates were imaged on a Nikon Ti2 Eclipse fluorescent microscope for DAPI, GFP, and mCherry. Using Nikon Elements software for analysis, nuclei were segmented based on DAPI, and mean object intensity per cell for both TopGFP and the internal control (mCherry) was measured. To calculate the amount of Wnt activity, the mean intensity of TopGFP was divided by the mean intensity of mCherry per cell to normalize individual cells. Curves and EC50s were plotted and calculated using GraphPad Prism software.

### HCEC GI_50_ Assay

HCEC cells were seeded at 2,000 cells per well in 384-well black SCREENSTAR imaging microplates and allowed to adhere overnight. The following day, cells were stimulated to induce the WNT pathway using 10μM CHIR99021 (#SML1046, Sigma) or 8 nM WNT3A (#5036-WN, R&D). Simultaneously, DYR compounds were delivered in a dose-response using a Tecan d300e digital dispenser ranging from 0μM to 30μM concentrations. Cells were incubated for 24 hours before fixing for 30 minutes with 4% paraformaldehyde/sucrose solution. Cells were permeabilized with 0.1% triton-x in PBS for 10 minutes and stained for DAPI for 30 minutes. Plates were imaged on a Nikon Ti2 Eclipse fluorescent microscope for DAPI. Using Nikon Elements software for analysis, nuclei were segmented based on DAPI, and nuclei counts were extracted and plotted using GraphPad Prism software.

### RNA isolation and quantitative real-time PCR

Total RNA was isolated from cells using the PureLink™ RNA Mini Kit and cDNA was synthesized using RevertAid First Strand cDNA Synthesis Kit for PCR with reverse transcription (RT–PCR). qRT-PCR analysis was performed using the SYBR™ Green PCR Master Mix on QuantStudio 7 Real-Time PCR system (Applied Biosystems). Data were normalized to corresponding GAPDH levels. Primer details in Materials table.

### Statistics and data presentation

Details of all statistical tests and multiple comparisons used to derive p value has been detailed in Figure Legends. All experiments were repeated 2-3 times with multiple technical replicates to be eligible for the indicated statistical analyses, and representative image has been shown. All results are presented as mean ± SD unless otherwise mentioned. For animal studies, Mead’s resource equation was used to predetermine sample size. Mice bearing parental tumours were randomized into two or three groups prior to vehicle or drug treatment. The investigators were not blinded to allocation during experiments but were blinded to predetermined end-point assessment. Data were analyzed using Graphpad Prism 10 statistical package.

## Materials Table

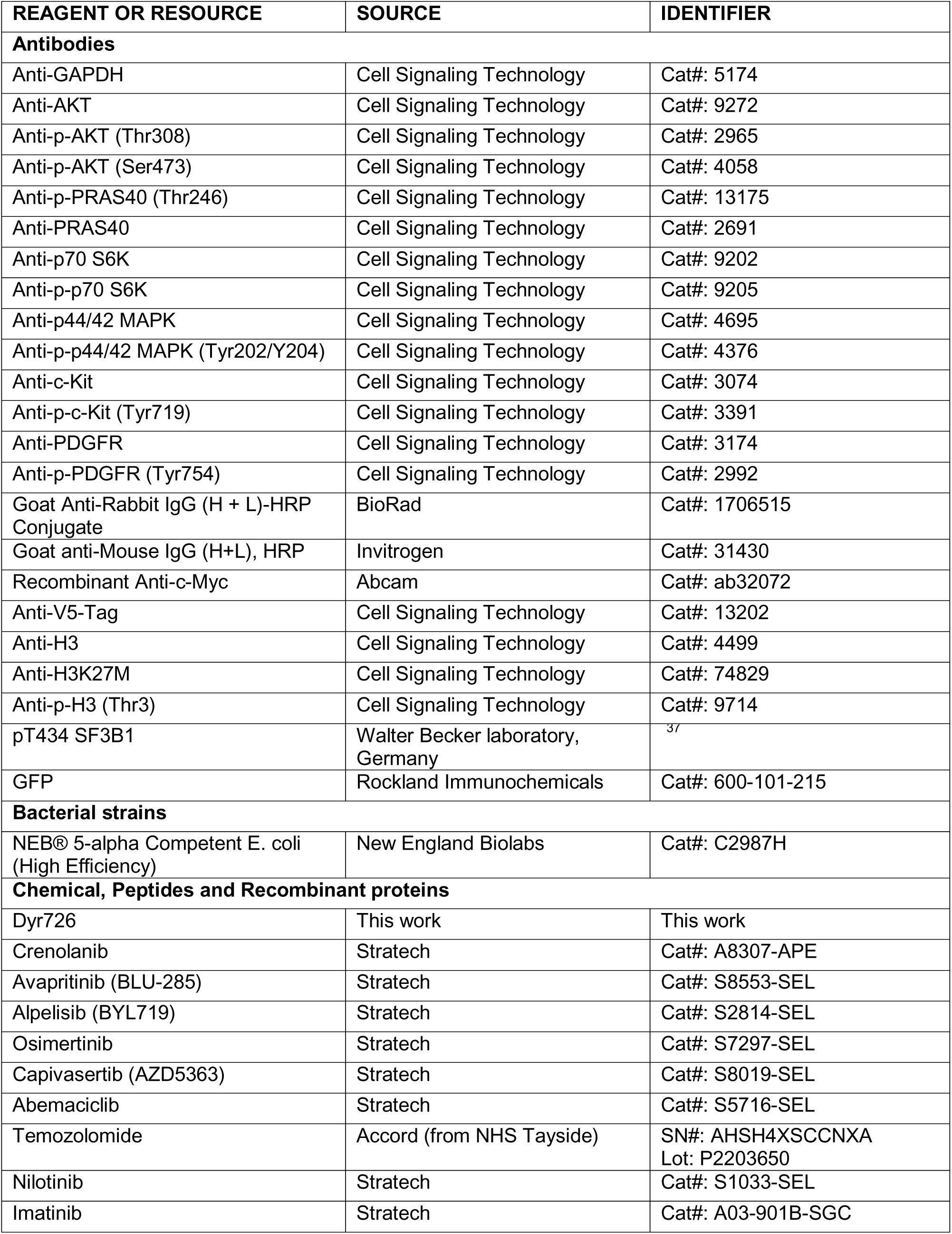
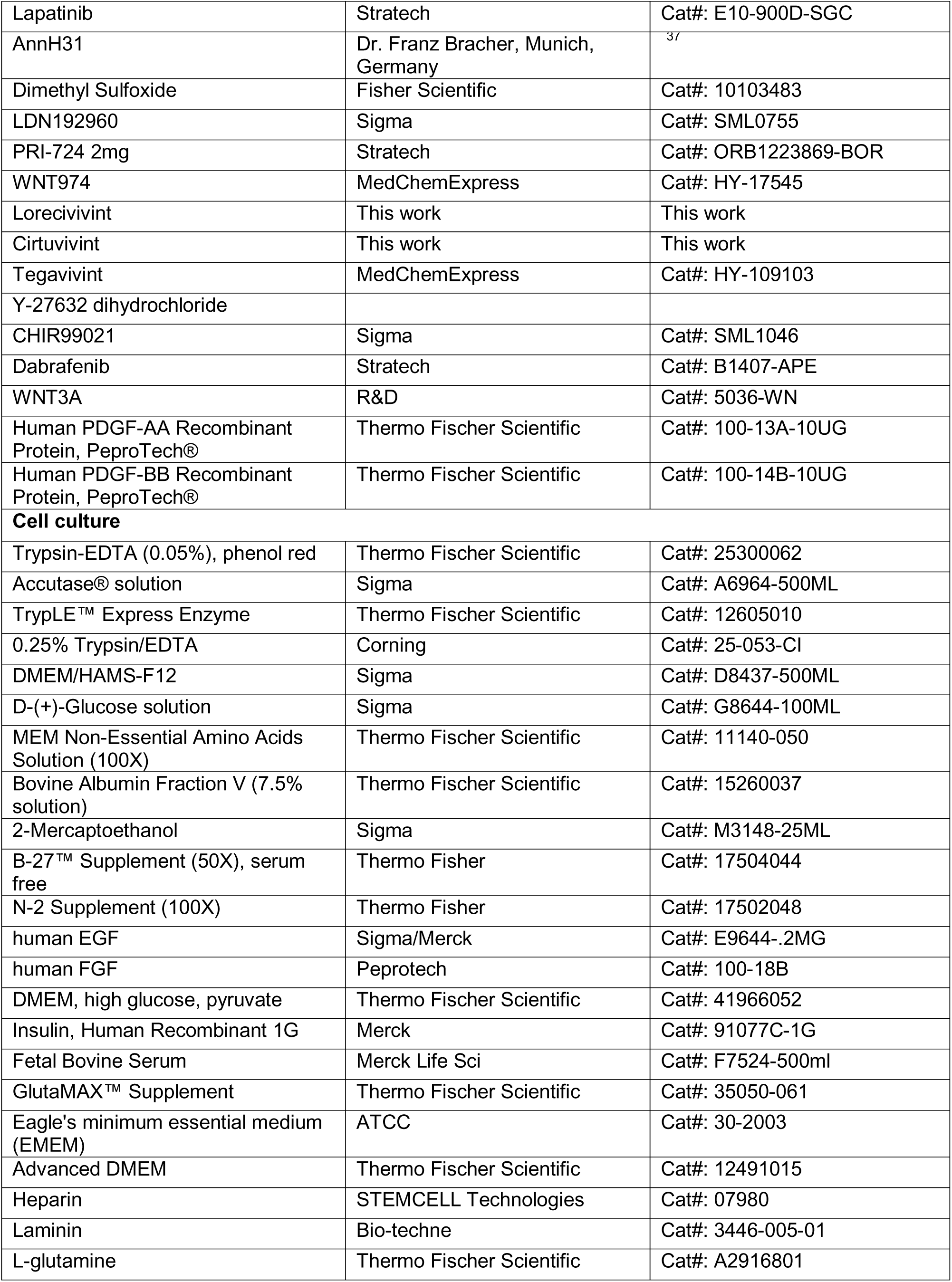
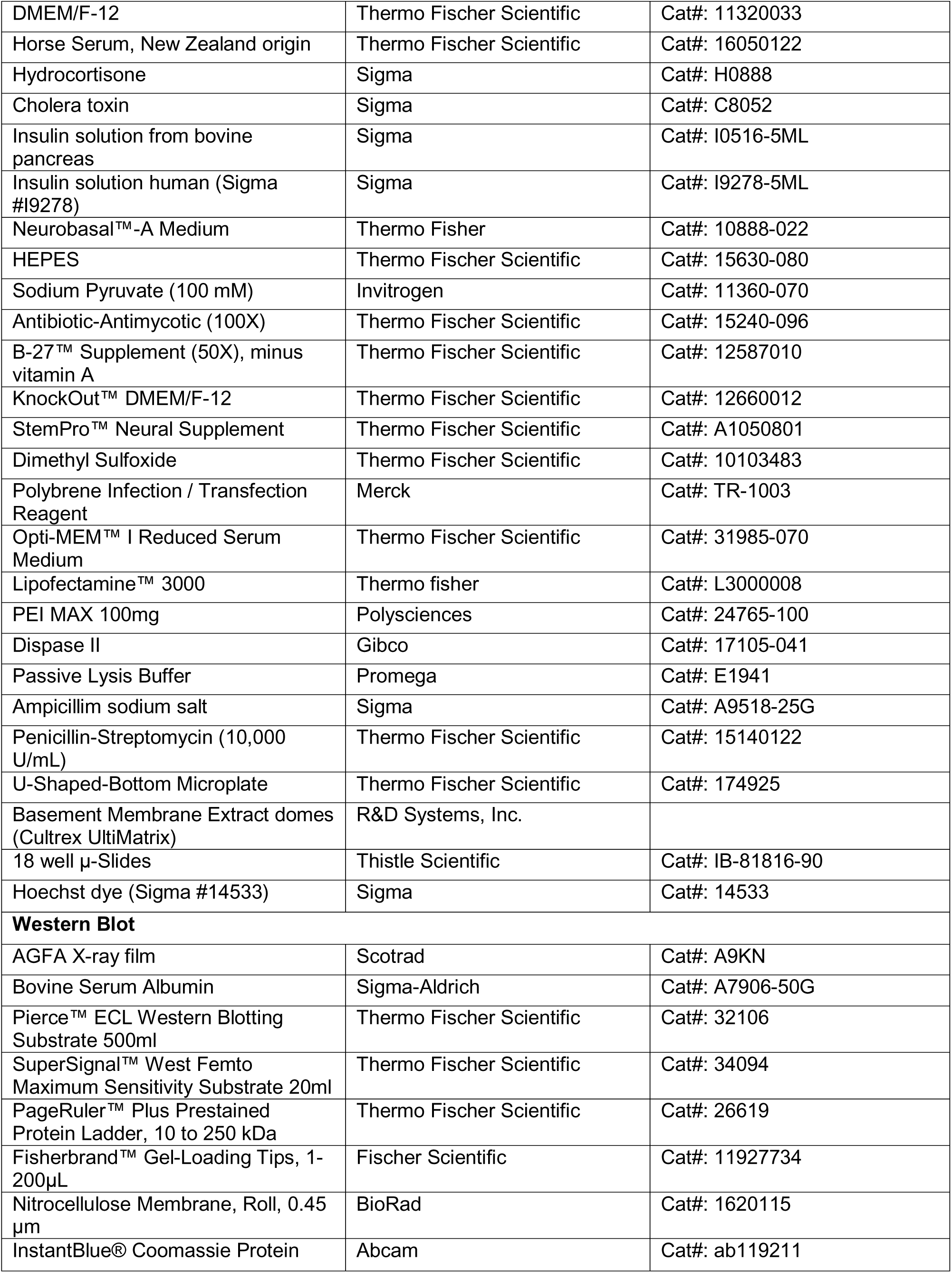
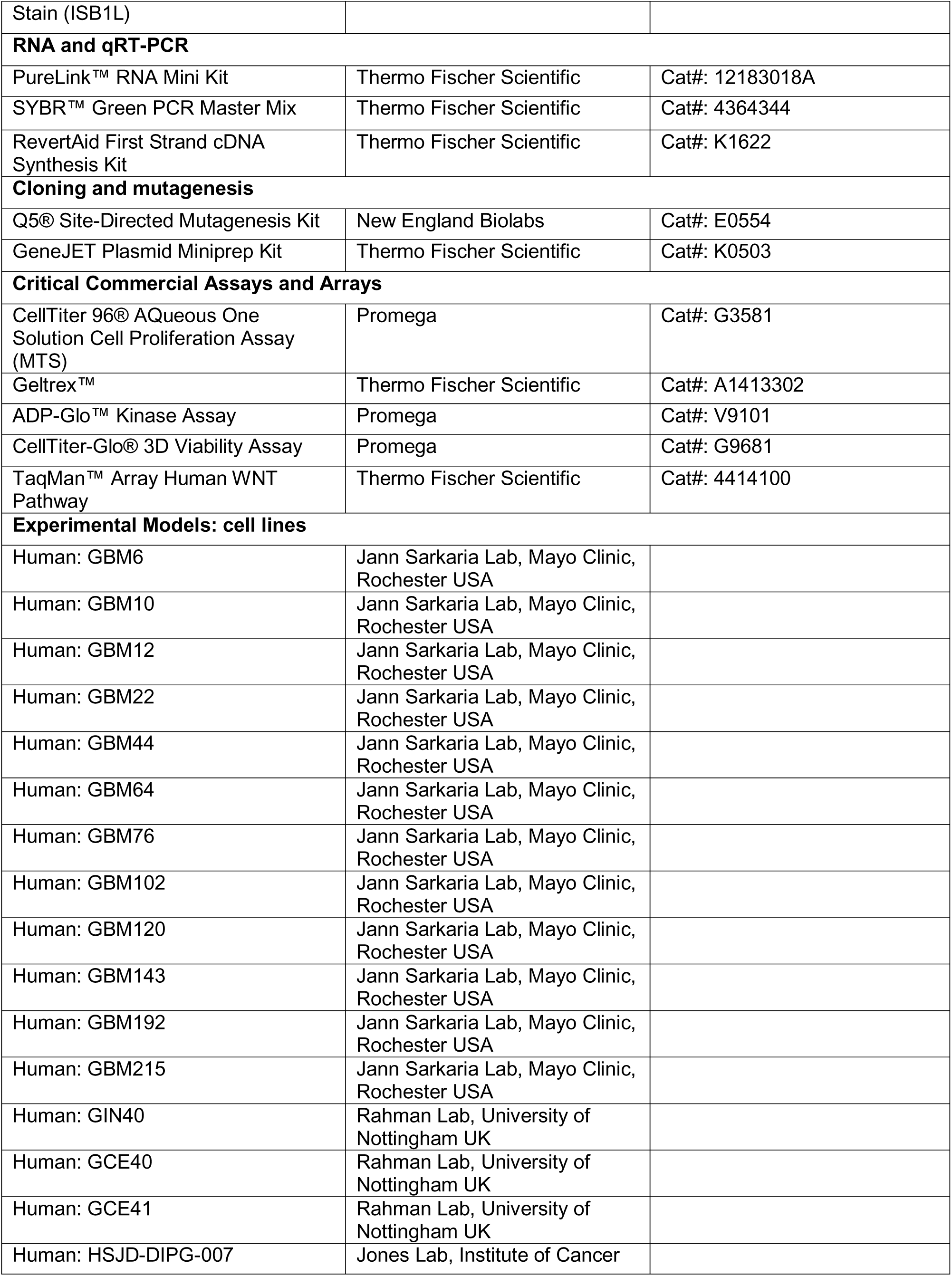
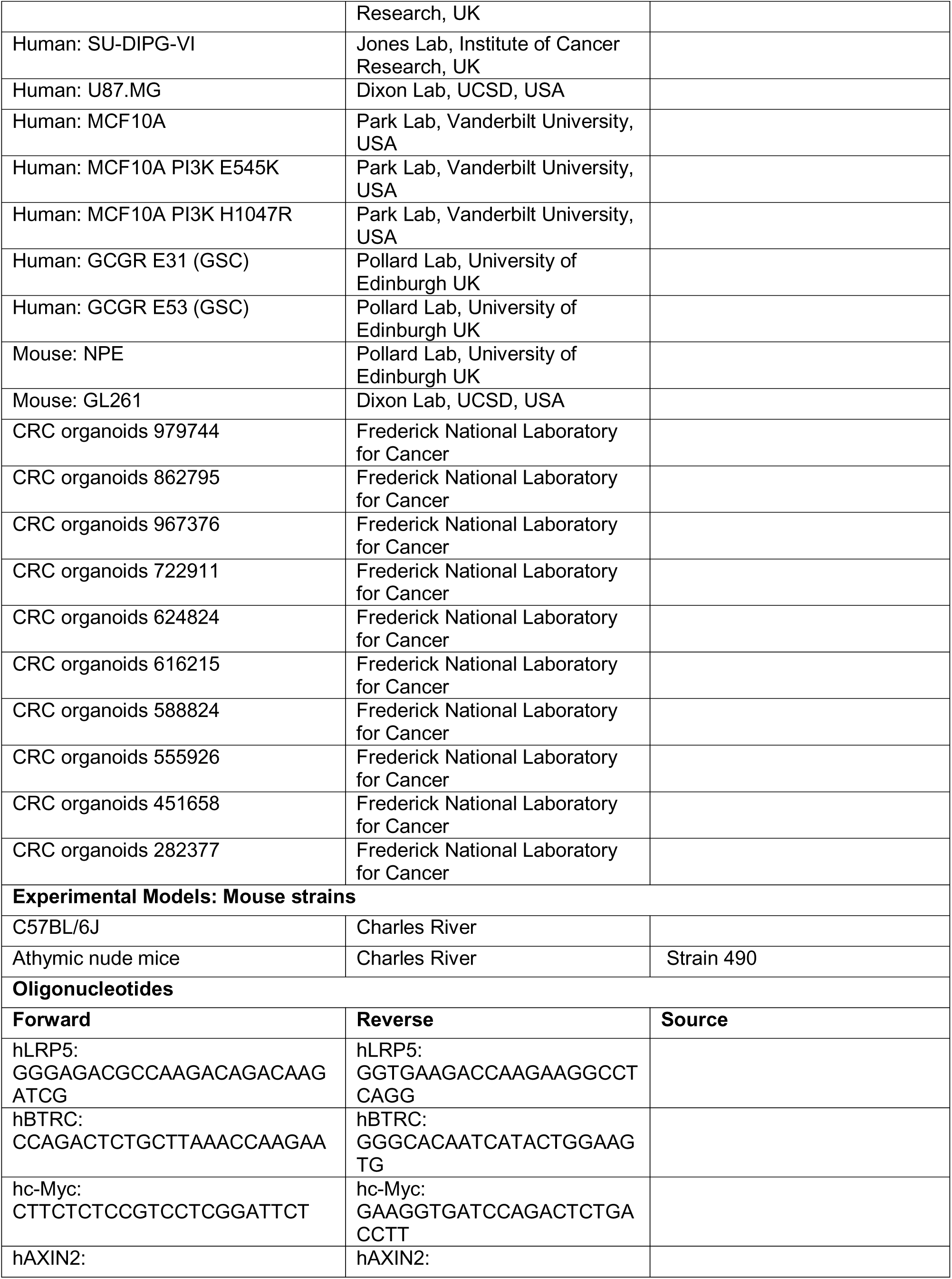
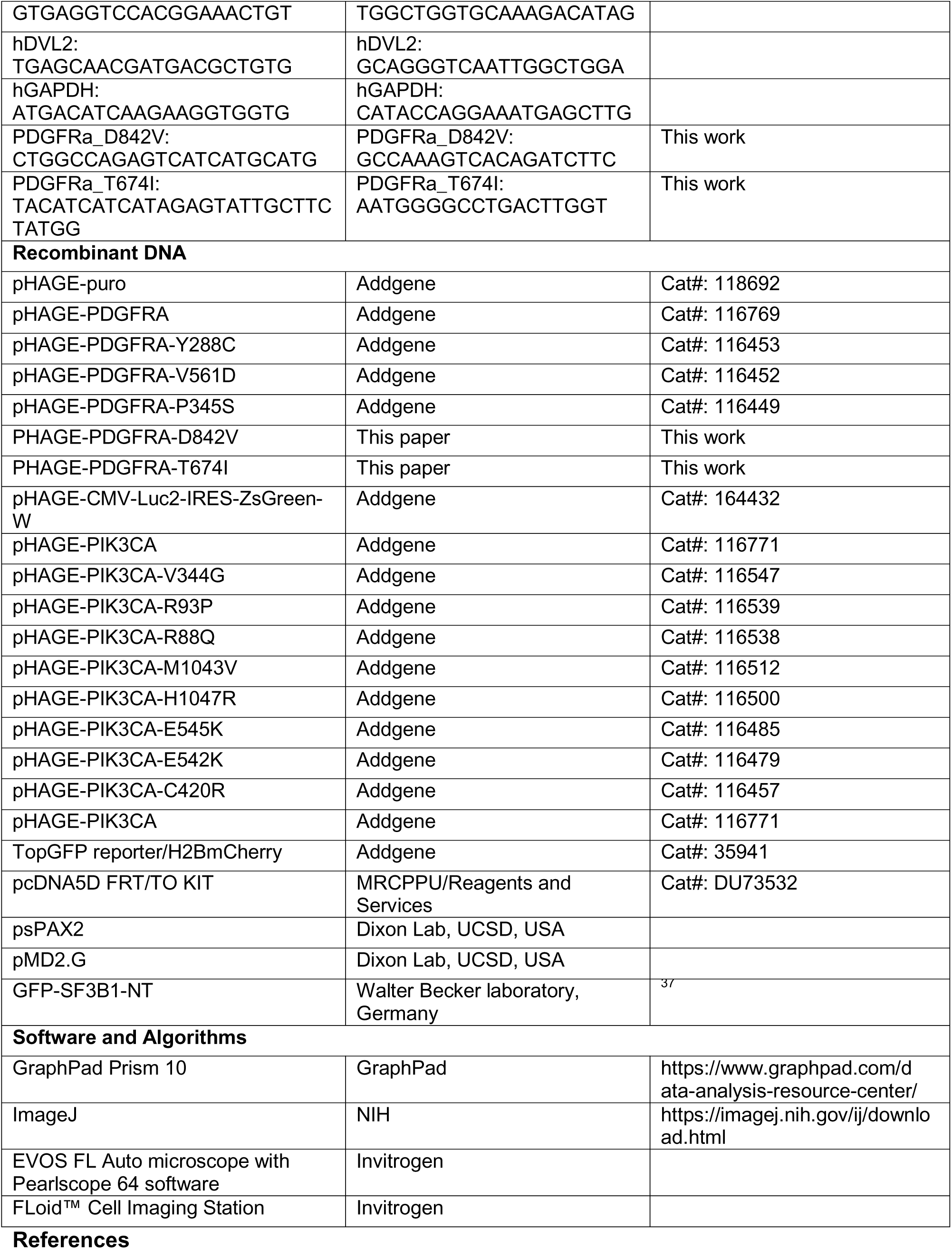

